# Rapid Changes in Transcription During a Feast-Famine Event

**DOI:** 10.64898/2026.02.04.703792

**Authors:** Paul Dijkstra, Bruce A. Hungate, Jennifer Pett-Ridge, Steven J. Blazewicz, Javier A. Ceja-Navarro, Ember M. Morrissey, Peter F. Chuckran, Egbert Schwartz

## Abstract

**Abstract:** Soil microbes have sophisticated mechanisms to detect and respond to short pulses of C inputs, often involving changes in gene-expression. We studied gene transcription in a soil microbial community before, and 8, 24, and 48h after glucose addition (0.7 mg C g^-1^ dry soil) to understand how microbes react to periods of substrate excess and subsequent starvation.

The relative transcript abundance of genes associated with energy metabolism and biosynthesis of amino acids, lipids, nucleotides, and cell wall components increased 8h after glucose addition. By 24 and 48h, the abundances of these transcripts reversed. Transcript abundance for genes associated with degradation of lipids, nucleotides, and (hetero)cyclic hydrocarbons decreased at 8h, but increased 24 and 48h after glucose addition. Simultaneously with a rise in transcripts for energy production and biosynthesis at 8h, transcription of regulatory genes for the exponential growth phase and ribosome assembly and maturation increased. In contrast, at 24 and 48h, transcript abundance for genes associated with ribosomal hibernation, sporulation, and regulation of the stationary phase increased, while transcripts for regulators for the exponential phase, and ribosome activation decreased. Based on changes in transcript abundance of phosphoenolpyruvate carboxylase and pyruvate carboxylase, it appeared that 8h after glucose addition glycolytic activity was high, however, gluconeogenesis returned at 24 and 48h. High levels of transcripts for *nrtC-ntrB* indicated N limitation 8 and 24h after glucose addition. Transcripts associated with Type VI Secretion Systems increased 24 and 48h after start of the experiment, suggesting a short lag between primary consumers and predatory bacteria. These results illustrate how metatranscriptome analysis can be used to study the ecophysiology of soil microbes providing details on the timing of exponential and stationary phase processes, coordination between anabolism and catabolism, and emerging nutrient limitations in natural soil communities.

**Research Highlights:**

1. We studied gene transcription of a soil microbial community after glucose addition
2. Transcript abundances for biosynthesis and energy production initially increased, while those for degradation decreased
3. Transcripts of regulators and sporulation genes indicated start of stationary phase at 24h
4. Nitrogen limitation induced transcription of nitrogen stress genes

## 1. INTRODUCTION

Soil microbes are important drivers of C and nutrient cycling and soil organic matter formation (Cotrufo et al., 2013, 2015; Lehmann and Kleber, 2015; Liang et al., 2017, 2020). However, we have limited knowledge of changes in microbial ecophysiology and growth, and the underlying processes of biosynthesis and energy production, in response to C inputs. Soil microbes experience repeated feast-famine events with short periods of growth and high activity separated by long periods of starvation (Lennon and Jones, 2011; Hobbie and Hobbie, 2013; Kuzyakov and Blagodatskaya, 2015). Much of the research on feast-famine events in soil communities has focused on respiration, microbial biomass dynamics (Anderson and Domsch, 2010), C and N cycling, and microbial C use efficiency (Reischke et al., 2015; Kallenbach et al., 2016; Hagerty et al., 2018; Manzoni et al., 2018; Geyer et al., 2019). However, it has proven difficult to directly determine the ecophysiological mechanisms underlying feast-famine responses.

Feast-famine transitions have been studied extensively in pure culture experiments (Brown et al., 2016; Jaishankar and Srivastava, 2017; Dworkin, 2023). Here, microbial activity is often separated into lag, exponential, and stationary phase. Rapid transitions in gene-expression occur between resource excess (exponential phase) and resource limitation (stationary phase), lasting only a few hours. These transitions are accompanied by large changes in transcription (e.g., Blencke et al., 2003; Koburger et al., 2005; Lemuth et al., 2008), mediated by signaling molecules (Valentini and Filloux, 2016; Irving et al., 2021) and master regulators of transcription (sigma factors, transcription factors and regulators, and DNA-binding proteins; Sharma and Chatterji, 2010; Paget, 2015). Accompanying the transition from growth to stationary phase are reductions in substrate consumption (Koburger et al., 2005), biosynthesis (Chang et al., 2002; Voigt et al., 2007), metabolic activity, increased utilization of alternative substrates (Wolfe, 2005; Voigt et al., 2007), alterations in the central C metabolic network, and transitions from glycolysis to gluconeogenesis (Lemuth et al., 2008; Handtke et al., 2018). The effects of starvation on amino acid biosynthesis are variable (Chang et al., 2002; Eymann et al., 2002; Voigt et al., 2007; Irving et al., 2021).

Here, we analyzed the transcriptional response of a soil microbial community to a sudden increase in glucose (simulating rhizosphere exudation) with the goal to improve our understanding of the microbial ecophysiology in soil ecosystems. Glucose addition experiments have a long history in soil ecology (Anderson and Domsch, 1985a, b; Burns, 2010) and have fueled influential concepts regarding microbial activity, substrate quality and quantity, and efficiency of substrate use. A previous publication based on this experiment (Chuckran et al., 2021) focused on transcription of genes of the N cycle and reported large changes in transcript abundances for ammonium and nitrate transporters, nitrate and nitrite reduction, ammonium assimilation via glutamate synthase and glutamine synthetase (GS-GOGAT), and regulation. Here, we describe the ecophysiological processes underlying the C cycle response during a feast-famine event, focusing on the processes underlying biomass production (biosynthesis, degradation, energy production, central C metabolism and gluconeogenesis) and regulators of cellular activity (exponential and stationary phase, ribosomal activation and hibernation), DNA replication, cell division, sporulation, and protein secretion. We use the terminology of primary vs. secondary consumers, with primary consumption defined as direct consumption of the added substrate, and secondary consumption as the utilization of products of primary consumers. Secondary consumption includes predation, scavenging, and cannibalism (González-Pastor, 2011; Gophna, 2022), but also utilization of waste products, extracellular enzymes, and biofilm. We explored the use of metatranscriptomes in intact soil communities for understanding the dynamics of growth and stationary phase processes, their regulation, primary and secondary consumption, and nutrient limitation during a feast-famine event.

### 2. MATERIALS AND METHODS

Soil (0-10 cm depth) was collected from a long-term crop rotation experiment (Pena-Yewtukhiw et al., 2017; Walkup et al., 2020) at the West Virginia University Certified Organic Farm (Morgantown, WV, USA; 39.647502°N, 79.93691°W). Soils in the experimental area are characterized as fine-loamy, mixed, active, mesic Ultic Hapludalf, fine-loamy, mixed, superactive, mesic Oxyaquic Hapludalf, and fine-silty, mixed, semiactive, mesic Typic Fragiudult. Soil was shipped to Flagstaff, Arizona, homogenized and sieved (2 mm mesh). Subsamples were preincubated in Mason jars (30 g soil) for two weeks at room temperature (23°C), after which each sample was mixed with 0.7 mg of glucose-C g^-1^ dry soil while raising soil moisture to 80% of field capacity. Soil for DNA and RNA extraction was collected just before and 8, 24, 48 hours after glucose addition (n=4 per timepoint), immediately frozen in liquid nitrogen, and stored at −80 °C. On parallel samples, respiration was measured using a Licor 6262 (Li-Cor Industries, Omaha, NE, USA) and nitrate and ammonia concentrations were determined using a SmartChem 200 discrete analyzer (Westco Scientific Instruments, Brookfield, CT, USA). Dissolved organic C (DOC) was determined in 0.5 M K_2_SO_4_ extracts using a TOC-L (Shimadzu Corporation, Kyoto, Japan). Further details of this experiment and results of measurements of respiration, dissolved inorganic N, DOC, MBC, and MBN were described in Chuckran et al. (2020, 2021; Supplementary Fig. 1).

### 2.1 DNA and RNA extraction and data processing

RNeasy PowerSoil total RNA and RNeasy PowerSoil DNA elution kits (Qiagen) were used to extract RNA and DNA according to the manufacturer’s instructions. RNA extracts were treated with an RNase-free DNase (Qiagen) to remove any DNA. Nucleic acid concentrations were quantified using a Qubit fluorometer (Invitrogen, Carlsbad, CA, USA) and purity was determined using a NanoDrop ND-1000 spectrophotometer (Nanodrop Technologies, Wilmington, DE, USA).

High-quality RNA samples were ribo-depleted (to remove most of ribosomal RNA; He et al., 2010) and sequenced by the Joint Genome Institute (JGI) using an Illumina NovaSeq platform (Illumina Inc., San Diego, CA, USA; Nordberg et al. (2013); 64-440 Mbp per assembled metatranscriptome). Details on library preparation, sequencing protocols, and samples are available in Chuckran et al. (2020). Reads were filtered using BBTools v38 (Mukherjee et al., 2019), assembly of metagenomes was done using SPAdes version 3.13.0 (Bankevich et al., 2012), and metatranscriptomes were assembled using MEGAHIT v1.1.2 (Li et al., 2015). Three samples were rejected during sequencing preparation for quality reasons, resulting in 13 metagenomes (n=3-4 per time point) and 16 metatranscriptomes (n=4; Chuckran et al. 2020). Read mapping was done using BBmap (Mukherjee et al., 2019) and annotation of assembled contigs was done using the IMG Annotation Pipeline v5.0.1 (Huntemann et al., 2016; Chen et al., 2019).

For this study, we downloaded KEGG (Kyoto Encyclopedia of Genes and Genomes; Kanehisa and Goto, 2000) annotated gene and transcript counts from the IMG website (Chuckran et al. 2020). We normalized read counts across samples and timepoints using DESeq2 (Love et al., 2014). Using pre-existing KEGG pathways and modules as a guide (https://www.genome.jp/kegg/), we manually curated metabolic pathways with the goal of minimizing overlap (Supplementary Table 1). Statistical analysis of (summed) gene and transcript abundances was done using ANOVA with the assumption of equal variance. If deviations from equal variance were significant (Levene’s test, R package ‘car’; Fox and Weisberg, 2019), ANOVA with unequal variance was used. If significant deviations from normality (Shapiro’s test) were observed, we used a non-parametric Kruskal-Wallis test for significance. Means significance testing was done with Tukey’s Honest Significant Difference test for Anova and Dunn’s Test for Multiple Comparisons in case of deviations from normality. Statistical tests were done using R (R core Team, 2023). We made 134 statistical tests of changes in gene and transcript abundances over time, respectively. The p-values were determined using a family-wise false discovery rate of 5% using the P.adjust command in R with method “BH” (Benjamini-Hochberg). Means per time point and results of statistical testing are provided in Supplementary Tables 3 and 4. The abundances of genes and transcripts are reported in abundance per million genes or transcripts and as trends over time plotted as:

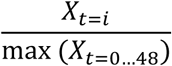

with X_t=i_ as the transcript or gene abundance at time *i* (0, 8, 24 or 48h). With this transformation, the maximum value of transcript or gene abundance over time always equaled 1.0. This transformation facilitates comparisons of temporal variation of genes and transcripts with very different absolute abundances. Changes in gene abundances over time in response to glucose addition for the studied processes were not significant (Supplementary Table 3, Supplementary Figs. 8-19).

### 2.2 Data availability

Raw sequence reads, assembled contigs, taxonomic and functional annotations are available under GOLD project identifier Gs0135756 (https://genome.jgi.doe.gov/portal/). IMG identification numbers, sample names, treatments, and sample descriptions are available in Chuckran et al. (2020).

## 3. RESULTS

We analyzed gene and transcript abundances for 1938 genes (Supplementary Table 1) over 48 hours following a glucose addition. These genes represented 19% of total number of genes and 22% of genes with transcripts (Supplementary Table 2). The summed abundance of these genes was about 35% of the total gene abundance and did not change significantly over time (Supplementary Tables 2 and 3, Supplementary Figs. S8-S19). However, the summed transcript abundance for these genes, as a proportion of the total transcript pool, increased from 33% before to more than 56% 8h after glucose addition (Supplementary Table 2).

### 3.1 Energy Production

Most organic C substrates are processed via the central C metabolic network (Chen et al., 2016; Dolan and Welch, 2018; Schink et al., 2022). We hypothesized that after glucose addition, transcript abundances for the processes of the central C metabolic network would increase, reflecting increased demand for energy and biosynthetic precursors. Relative transcript abundances for the four main pathways of the central C metabolic network (glycolysis, TCA cycle, pentose phosphate pathway and Entner-Doudoroff pathway) increased significantly 8h after glucose addition but decreased again significantly at 24 and 48h (Fig. 1A, Supplementary Table 4). Most of the transcripts were associated with glycolysis and TCA cycle (Supplementary Fig. 2A). Transcript abundances for the glyoxylate bypass were significantly higher 24h and 48h after glucose addition (Fig. 1A, Supplementary Table 4).

**Fig. 1.**
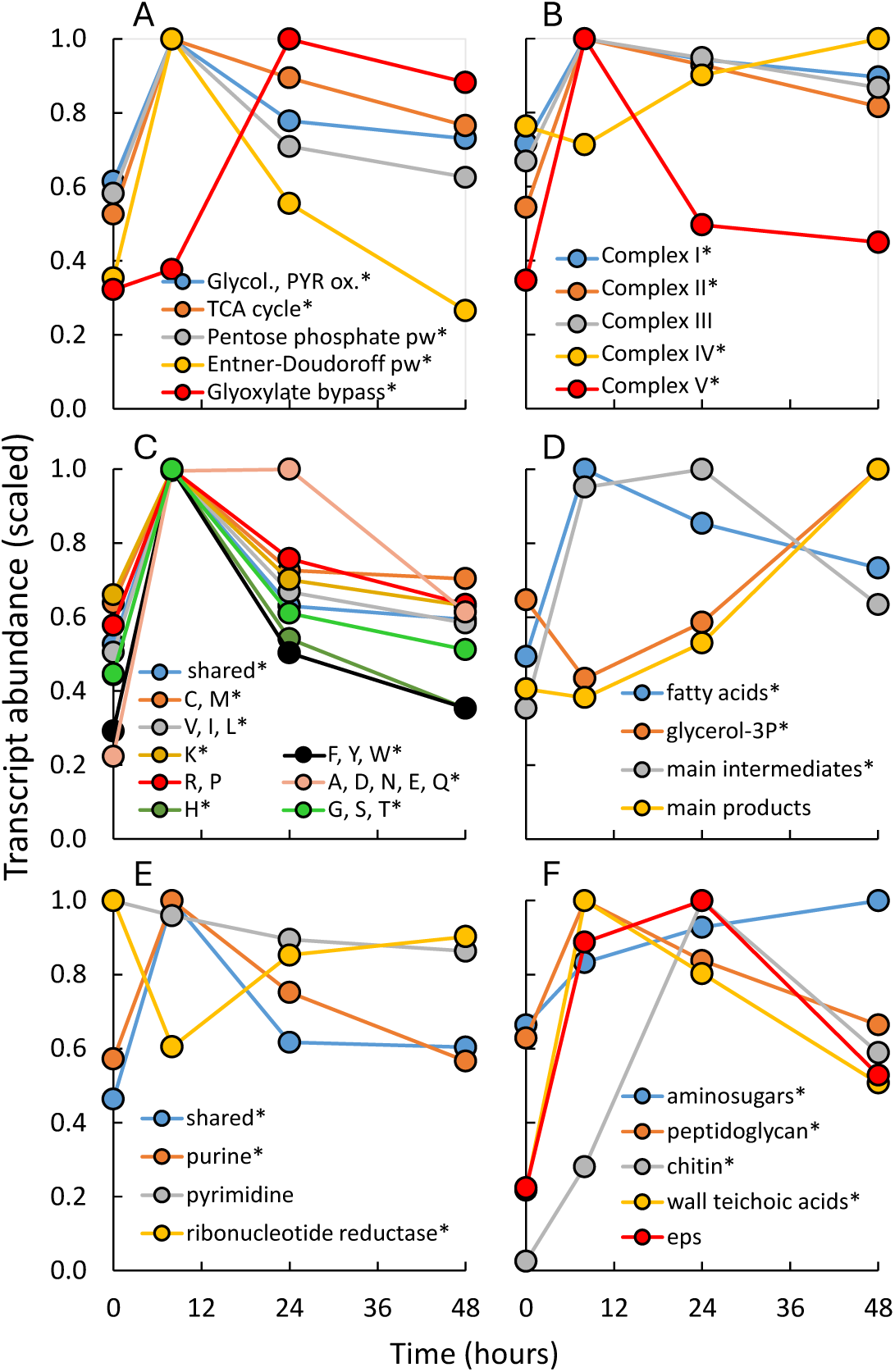
Effects of glucose addition on transcript abundances (scaled relative to maximum abundance) for genes associated with (A) central C metabolic network, (B) electron transport chain, biosynthesis of (C) amino acids, (D) lipids, (E) nucleotides, and (F) cell wall compounds and exopolysaccharides. * indicates statistically significant differences across time (n=4; results from multiple mean tests in Supplementary Table 4). Glycol., PYR ox. stands for glycolysis and pyruvate oxidation, pw. for pathway, and EPS for exopolysaccharides. Amino acid biosynthesis pathways are represented by their one-letter amino acid code. Information on gene and transcript identity, abundances and additional statistical information is available in Supplementary Information.

NADH produced via the central C metabolic network processes is used to generate a proton or sodium ion gradient via the electron transport chain (ETC; Sazanov, 2015). Transcript abundances of ETC Complex I and II were significantly higher after glucose addition and decreased slightly (NS) at 24 and 48h (Fig. 1B, Supplementary Fig. 2B, Supplementary Table 4). Transcript abundances for Complex III showed a similar but not-significant trend. Transcripts for genes of ETC Complex IV gradually but significantly increased over time, while transcripts for ETC Complex V, the F-type ATP-synthase, showed a sharp and significant increase followed by a steep and significant decline at 24 and 48h (Fig. 1B).

### 3.2 Biosynthesis

We hypothesized that transcripts associated with biosynthesis pathways would increase after glucose addition. The number of transcripts for proteinogenic amino acid biosynthesis significantly increased 8h after glucose addition but dropped (P<0.05) after 24 and 48h (Fig. 1C, Supplementary Fig. 2C, Supplementary Table 4). All major amino acid biosynthesis pathways showed similar patterns, except for alanine, aspartate, asparagine, glutamate and glutamine biosynthesis. Before glucose addition, transcripts for the biosynthesis of this group of amino acids made up 23% of the transcript pool (Supplementary Fig. 2C). This proportion increased to 40-50% after glucose was added to the soil, with transcripts for one enzyme, GlnA (glutamine synthetase) making up 16% before glucose addition and 30-40% thereafter.

Biosynthesis of lipids is an essential function in all organisms (Sohlenkamp and Geiger, 2015; Maurya et al., 2019; Allen and Martinez, 2020). The lipid composition of the membrane affects important processes such as protein export, respiration, and adaptation to temperature and osmotic stress (Arias-Cartin et al., 2012). After adding glucose, transcript abundance for fatty acid production (peak 8h after glucose addition) and main intermediates (peak at 8 and 24h) significantly increased (Fig. 1D, Supplementary Fig. 2D, Supplementary Table 4), while transcripts associated with the main lipid products showed no significant change. The transcript abundance for glycerol-3P biosynthesis gradually and significantly increased towards the end of the experiment. Transcripts for phosphatidylglycerol (peak 8h), triacylglycerol, and glycolipids (peak 24h) increased significantly after glucose addition, while those for phosphatidylcholine significantly decreased at 8h before recovering towards the end of the experiment (Supplementary Table 4).

Transcript abundances for biosynthesis of purine nucleotides significantly increased at 8h but then significantly decreased 24 and 48h after glucose addition (Fig. 1E, Supplementary Fig. 2E, Supplementary Table 4) showing parallel patterns to transcripts for amino acids biosynthesis. In contrast, transcript abundances for pyrimidine biosynthesis did not change significantly and that associated with ribonucleotide reduction significantly decreased 8h after glucose addition.

Amino sugars are a major constituent of peptidoglycan and chitin (Barreteau et al., 2008; Lenardon et al., 2010). Transcripts for biosynthesis of amino sugars and peptidoglycan units significantly increased after glucose addition (Fig. 1F, Supplementary Fig. 2F). Transcripts for chitin synthase (*CHS1*) also significantly increased after glucose addition but reached a maximum at 24h. In Gram-positive bacteria, (wall) teichoic acids can make up 60% of the cell wall weight (Brown et al., 2013). Transcript abundance for teichoic acid biosynthesis rose sharply and significantly after glucose addition, peaking at 8h before (NS) declining at 24 and 48h (Fig. 1F, Supplementary Table 4).

Biofilm formation is essential for many soil microbes (McDougald et al., 2012). Transcript abundances for exopolysaccharide (EPS) production significantly increased 8 and 24h after glucose addition and (NS) decreased thereafter (Fig. 1F). The transcriptome of genes associated with biofilm polysaccharide biosynthesis significantly changed over time (Supplementary Table 4) with transcripts for vibriopolysaccharide and enterobacterial common antigen peaking 8h after glucose addition, while abundance for colonic acid, alginate, and succinoglycan peaked at 24 and for poly N-acetylglucosamine at 48h. Transcripts for bacterial cellulose biosynthesis, a component of biofilm in some bacterial species (Abidi et al., 2021; Acheson et al., 2021), increased significantly 8 and 24h after glucose addition (Supplementary Table 4).

Of transcripts associated with biosynthesis, those for amino acids made up about 50% before and 70% 8h after glucose addition, demonstrating the quantitative importance of amino acid biosynthesis in the cell’s C, nutrient, and energy budget (Supplementary Fig. 3A).

### 3.3 Degradation

Degradation of compounds other than carbohydrates provides a source of C, N, and energy and is part of regular turnover of cellular compounds. We hypothesized that degradative processes would transiently be reduced after glucose addition. Transcript abundances for degradation of amino acids did not significantly change in response to glucose addition (Fig. 2B, Supplementary Fig. 3B, Supplementary Table 4), although the individual pathways exhibited a variety of responses. There were significant decreases in transcript abundances for fatty acid, pyrimidine, and purine degradation 8h after glucose addition.

**Fig. 2.**
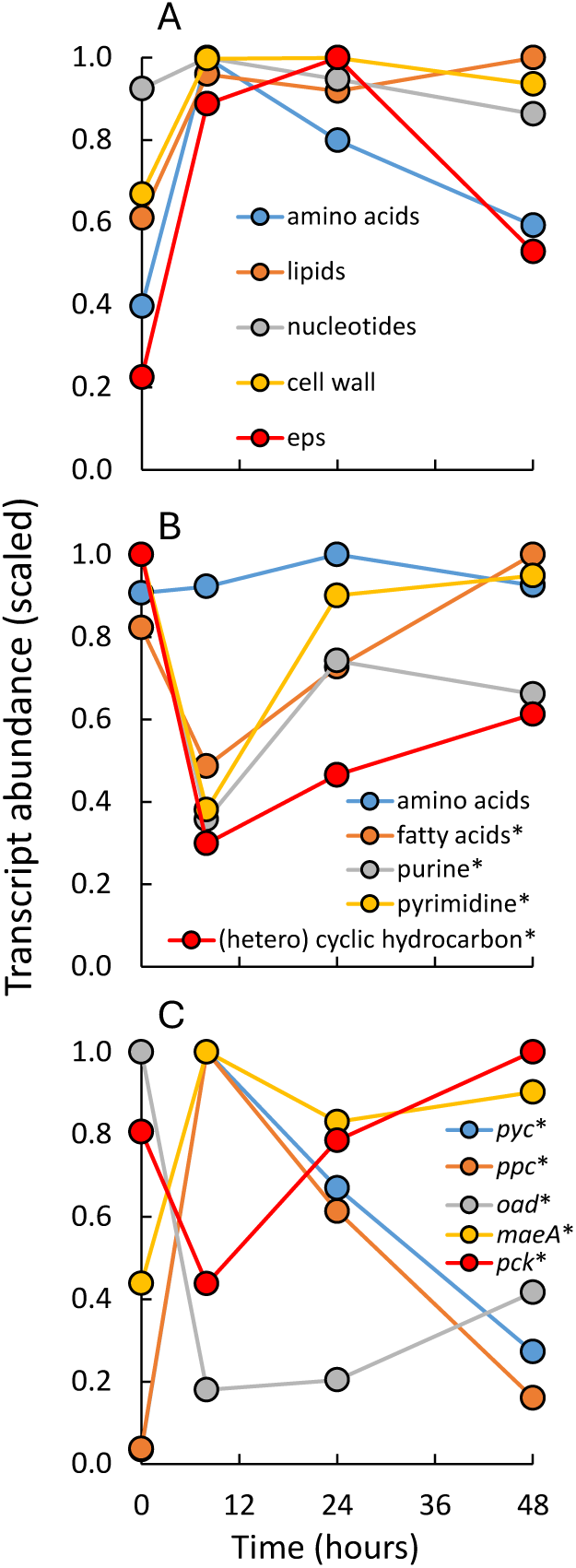
Effects of glucose addition on transcript abundances (scaled relative to maximum abundance) for genes associated with (A) biosynthesis of amino acids, lipids, nucleotides, cell walls and exopolysaccharides, (B) degradation of amino acids, lipids, nucleotides, and (hetero)cyclic hydrocarbons, and (C) genes for glycolytic (pyruvate carboxylase - *pyc*, phosphoenolpyruvate carboxylase - *ppc*) and gluconeogenic (oxaloacetate decarboxylase - *oad*, (oxaloacetate-decarboxylating) malate dehydrogenase - *maeA*, and phosphoenolpyruvate carboxykinase - *pck, pepck*) reactions between PEP, pyruvate and oxaloacetate and malate. * indicates statistically significant differences across time (n=4; results from multiple mean tests in Supplementary Table 4). Information on gene and transcript identity, abundances and additional statistical information is available in Supplementary Information.

Homocyclic and heterocyclic hydrocarbons, aromatic and non-aromatic, are generally viewed as recalcitrant compounds, and are thought to make up a substantial portion of soil and plant organic matter. Transcript abundances for genes associated with degradation of these compounds decreased significantly 8h after glucose addition but (NS) trended higher again at 24 and 48h (Fig. 2B, Supplementary Fig. 3B, Supplementary Table 4). Most of the genes and transcripts within this category were associated with benzoate and nitrobenzoate degradation pathways. Many of the genes are also involved in amino acid, nucleotide, and lipid degradation. Complicating interpretation further, some of these degradation pathways are linked to biosynthesis of secondary compounds (Craney et al., 2013). Most of the genes had low transcript abundances, however, cyclohexanone monooxygenase (*chnB*), gluconolactonase (*gnl*), carboxymethylenebutenolidase, and the large (*hyaB*) and small (*hyaA*) subunits of a hydrogenase were transcribed at high rates.

### 3.4 Substrate Use, Glycolysis, and Gluconeogenesis

Enzymes that drive the reactions between phosphoenolpyruvate, oxaloacetate, malate and pyruvate form an important control point in the central C metabolic network (Sauer and Eikmanns, 2005; Koendjbiharie et al., 2020), with reactions from malate and oxaloacetate to pyruvate and phosphoenolpyruvate stimulated in the absence of glucose (gluconeogenesis). We hypothesized that transcript abundances for these enzymes might differentiate between consumption of glucose and non-glucose substrates. Before glucose addition, transcripts for phosphoenolpyruvate carboxykinase (*pck, pepck*) were abundant (Supplementary Fig. 3C), suggesting initially high gluconeogenesis activity. Eight hours after adding glucose, transcript abundances for pyruvate carboxylase (*pyc*) and phosphoenolpyruvate carboxylase (*ppc*) significantly increased, consistent with glucose consumption and high glycolysis activity (Fig. 2C, Supplementary Fig. 3C, Supplementary Table 4). Transcript abundance of these glycolytic genes significantly declined again at 24 and 48h. Changes in transcript abundances for oxaloacetate decarboxylase (*oad*) and phosphoenolpyruvate carboxykinase (*pck*) showed the opposite pattern: significant decreases 8h and increases 24 and 48h after glucose addition. Surprisingly, transcript abundance for the glucogenic (oxaloacetate-decarboxylating) malate dehydrogenase (*maeA*) increased after glucose addition and remained high even after transcript abundance for pyruvate and phosphoenolpyruvate carboxylase decreased. The increased *maeA* transcript abundance may be related to the operation of the malate shunt, by-passing the PEP-pyruvate transformation by pyruvate phosphate dikinase under anaerobic conditions (Koendjbiharie et al., 2020). These results suggest that peak glucose utilization occurred 8h after glucose was added to the soil.

### 3.5 Resource Limitation after Glucose Addition

We hypothesized that after glucose addition, other nutrients would become limiting. We compared the transcript abundances for three groups of genes related to C, N, and P limitation (Schumacher et al., 2013; Shimizu, 2013; Santos-Beneit, 2015). The gene *crp* encodes a transcriptional regulator that controls C catabolic repression, the utilization of a preferred over less-preferred substrate (Ishizuka et al., 1993). Transcript abundances for *crp* significantly declined 8h after glucose addition, likely suppressed by the presence of glucose (Fig. 3A, Supplementary Fig. 4A, Supplementary Table 4). This suggests that glucose was not limiting microbial activity at this time. Transcript abundance for *phoB-phoR*, regulators of the P-stress response (Santos-Beneit, 2015), were moderately (NS) decreased 8h after glucose addition. However, transcript abundance for *ntrC-ntrB*, regulators of the N-stress response (Ludwig and Bryant, 2012; Schumacher et al., 2013; Brown et al., 2014) significantly increased (more than 15-fold): a sign of strong N limitation 8h and 24h after glucose addition. Transcription rates of genes associated with P-uptake and phosphatase activity (Martín and Liras, 2021) showed significant changes (Fig. 3B, Supplementary Fig. 4B, Supplementary Table 4), but to a much smaller degree than that for genes associated with N transport, uptake, assimilation, and regulation (Fig. 3C; Chuckran et al., 2021).

**Fig. 3.**
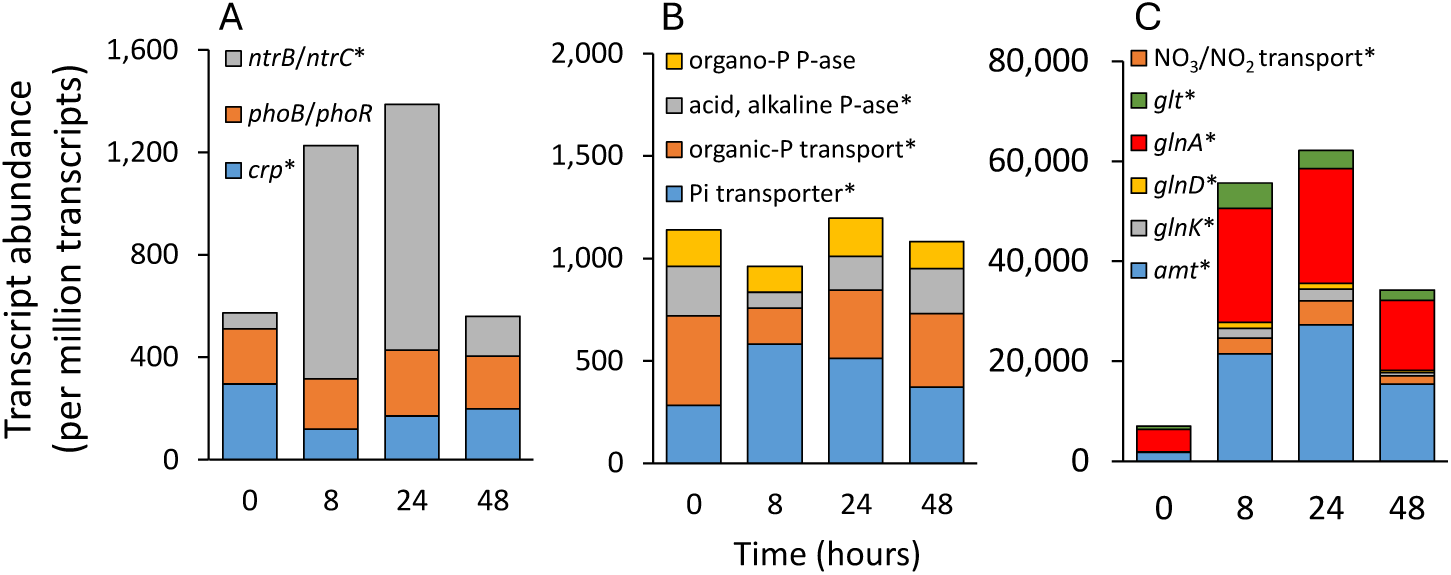
Effects of glucose addition on transcript abundance (transcripts per million transcripts) for (A) regulator genes associated with C (*crp*), P (*phoB-phoR*), and N stress (*ntrC-ntrB*), (B) P-transporters and phosphatases, and (C) ammonium (*amt*) and nitrate transporters, GS-GOGAT (*glnA*, *glt*) and regulatory genes (*glnK*, *glnD*). * indicates statistically significant differences across time (n=4; results from multiple mean tests in Supplementary Table 4). Information on gene and transcript identity, abundances and additional statistical information is available in Supplementary Information.

### 3.6 Transcription, Translation, DNA Replication, and Cell Division

We hypothesized that, after glucose addition, transcript abundances for transcription, translation, DNA replication, and cell division would increase. Transcript abundances for RNA polymerase and ribosomal proteins showed significant and large, although short-lived, increases in response to glucose addition (Fig. 4A, Supplementary Fig. 5A, Supplementary Table 4). In contrast, transcript abundance for DNA polymerase III did not significantly change over time.

**Fig. 4.**
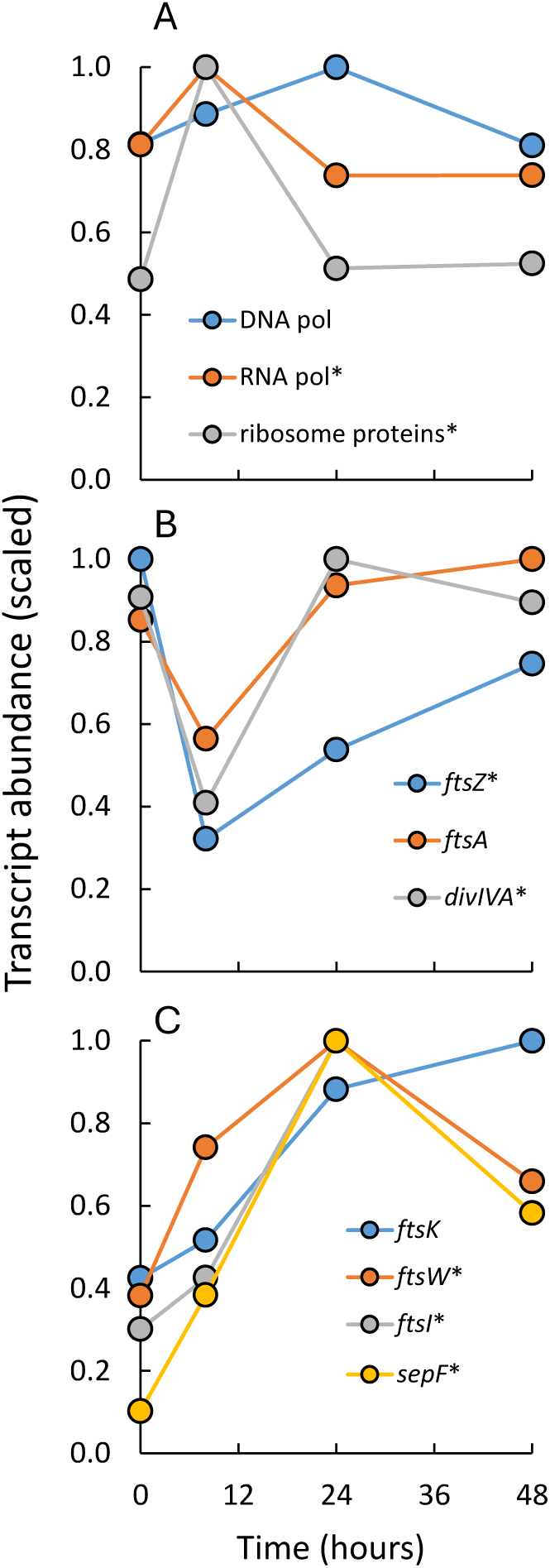
Effects of glucose addition on transcript abundances (scaled relative to maximum abundance) for (A) DNA polymerase III, RNA polymerase, and ribosomal proteins, (B) genes of early stage of cell division (*ftsZ, ftsA, divIVA)* and (C) mid-stage of cell division (*ftsK, ftsW, ftsI, and sepF*). * indicates statistically significant differences over time (n=4; results from multiple mean tests in Supplementary Table 4). Information on gene and transcript identity, abundances and additional statistical information is available in Supplementary Information.

Bacterial cell division starts with the formation of a Z-ring composed of FtsZ tubulin-like proteins at the site of the future cell wall (Haeusser and Margolin, 2016). These proteins form polymer filaments anchored to the membrane by, among others, FtsA proteins (Egan and Vollmer, 2013; Barrows and Goley 2021). For Gram-positive bacteria such as *Streptococcus*, DivIVA proteins plays an important role in cell division initiation and localization (Egan and Vollmer, 2013; Hammond et al., 2019). At mid-stage cell division, further enzyme machinery is recruited to the Z-ring, and a cell wall is formed. This additional machinery includes genes like *ftsK*, *ftsW*, *ftsI*, and other peptidoglycan synthesizing and modifying enzymes. Transcript abundances for *ftsZ* and *divIVA* genes, engaged in the early stage of cell division, were high before glucose addition, significantly reduced at 8h, and increased again at 24 and 48h (Fig. 4B; Supplementary Table 4). *FtsA* showed a similar but NS trend. Transcripts for FtsW, FtsI and SepF, mid-stage cell division proteins, showed peak abundances 24 or 48h after glucose addition (Fig. 4C). Changes in transcript abundances for *ftsA* and *ftsK* showed similar trends but were not significantly different over time (P<0.10).

### 3.7 Stationary Phase, Ribosome Hibernation, and Sporulation

Regulators of gene expression govern every life stage of microbial organisms. We hypothesized that transcripts for regulators could indicate transitions between the exponential and stationary phase. In *E. coli*, *fis* is a DNA-binding regulator expressed at the start of the exponential phase (Kahramanoglou et al., 2011; Brandi et al., 2016). Transcript abundance of *fis* in this experiment was high 8h after glucose addition, but significantly declined at 24 and 48h, suggesting that a portion of the microbial community transitioned to late exponential, early stationary phase by that time (Fig. 5A, Supplementary Fig. 6A, Supplementary Table 4). This conclusion is supported by the increased (P<0.06) transcript abundance for *hns* at 24h; this gene is characteristically expressed at the end of the exponential phase (Brandi et al., 2020).

**Fig. 5.**
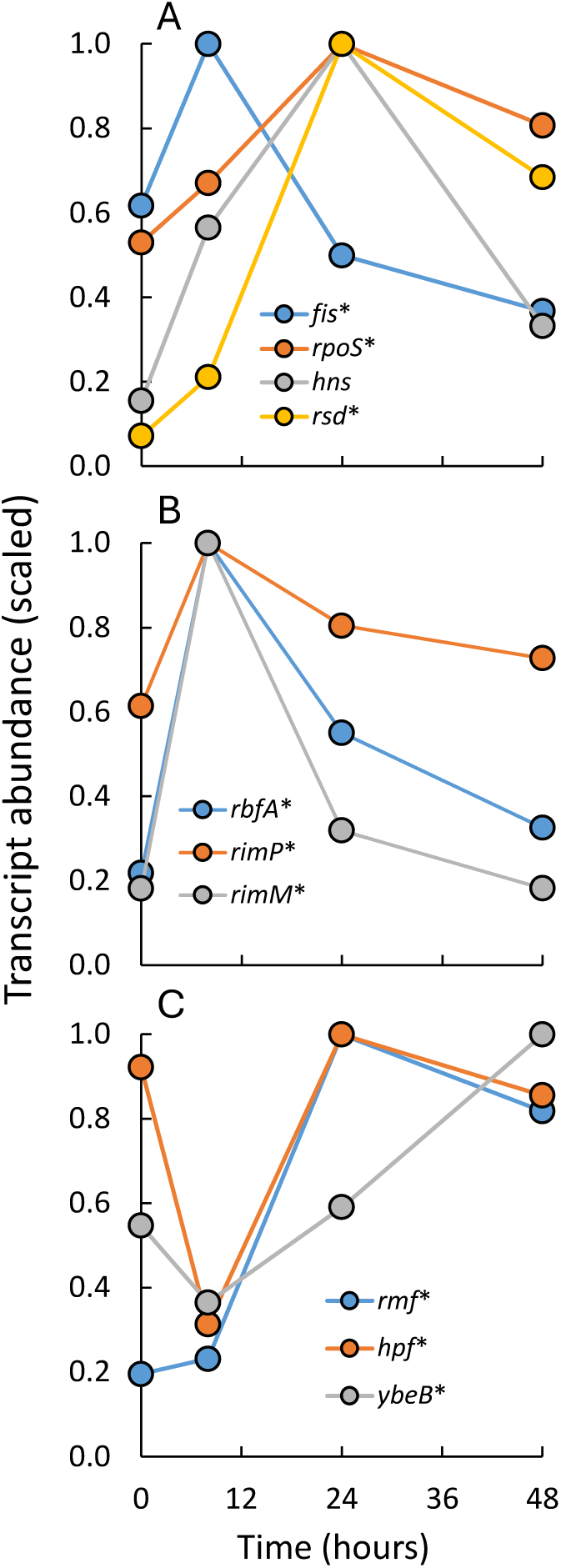
Effects of glucose addition on transcript abundances (scaled relative to maximum abundance) for (A) regulators of the early exponential growth phase (*fis*), late exponential growth phase (*hns*), and early stationary phase (*rpoS*, *rsd*), (B) ribosome activation (*rbfA*, *rimP*, and *rimM*) and (C) ribosome hibernation (*rmf*, *hpf*, *ybeB*). * indicates statistically significant differences over time (n=4; results from multiple mean tests in Supplementary Table 4). Information on gene and transcript identity, abundances and additional statistical information is available in Supplementary Information.

In response to starvation, regulators of the stationary phase are transcribed, including *rpoS* (Navarro-Llorens et al., 2010; Paget, 2015). Transcript abundance for *rpoS* significantly increased at 24h (Fig. 5A), suggesting that 24h after glucose addition, the stationary phase started. This is supported by the significant increase in transcript abundance for *rsd*, an anti-sigma D factor that binds to the constitutively expressed housekeeping sigma D (RpoD) protein (Hofmann et al., 2011), thereby reducing its role in regulation of gene-expression.

Rapid growth requires active and abundant ribosomes (Dennis et al., 2004). There was a large increase in transcripts for ribosomal proteins 8h after glucose addition (Fig. 4A, Supplementary Fig. 5A). The gene *rbfA* is required for ribosome subunit joining, maturation, and activity (Feaga et al., 2020; Maksimova et al., 2021). Transcript abundance for *rbfA* was significantly increased 8h after glucose addition followed by a significant decline at 24 and 48h (Fig. 5B; Supplementary Table 4), suggesting reduced ribosomal activity at that time. Similar transcriptional dynamics were observed for ribosome maturation factors *rimM* and *rimP* (Maksimova et al., 2022).

When feast turns into famine, ribosomal processes are among the first to be down-regulated (Matzov et al., 2019; Irving et al., 2021), reducing ribosome activity via ribosome hibernation, dimerization, and inactivation (Trösch and Willmund, 2019; Feaga et al., 2020). The process of dimerization is regulated by several genes, including *rmf* (Usachev et al., 2020), *hpf* (Feaga et al., 2020), and *ybeB*, a gene that blocks joining of ribosome subunits. In our experiment, transcript abundances for *rmf*, *hpf*, and *ybeB* were low 8h after glucose addition but significantly increased at 24 and 48h (Fig. 5C; Supplementary Table 4), suggesting ribosome dimerization and hibernation had started 24h after glucose addition.

In response to starvation and during the stationary phase, Bacillota species are known to start endospore formation (Riley et al., 2020). Before and 8h after glucose addition, abundances of transcripts associated with genes for endospore formation were low, but (significantly) increased 24 and 48h after glucose addition (Fig. 6A, Supplementary Fig. 7A, Supplementary Table 4).

**Fig. 6.**
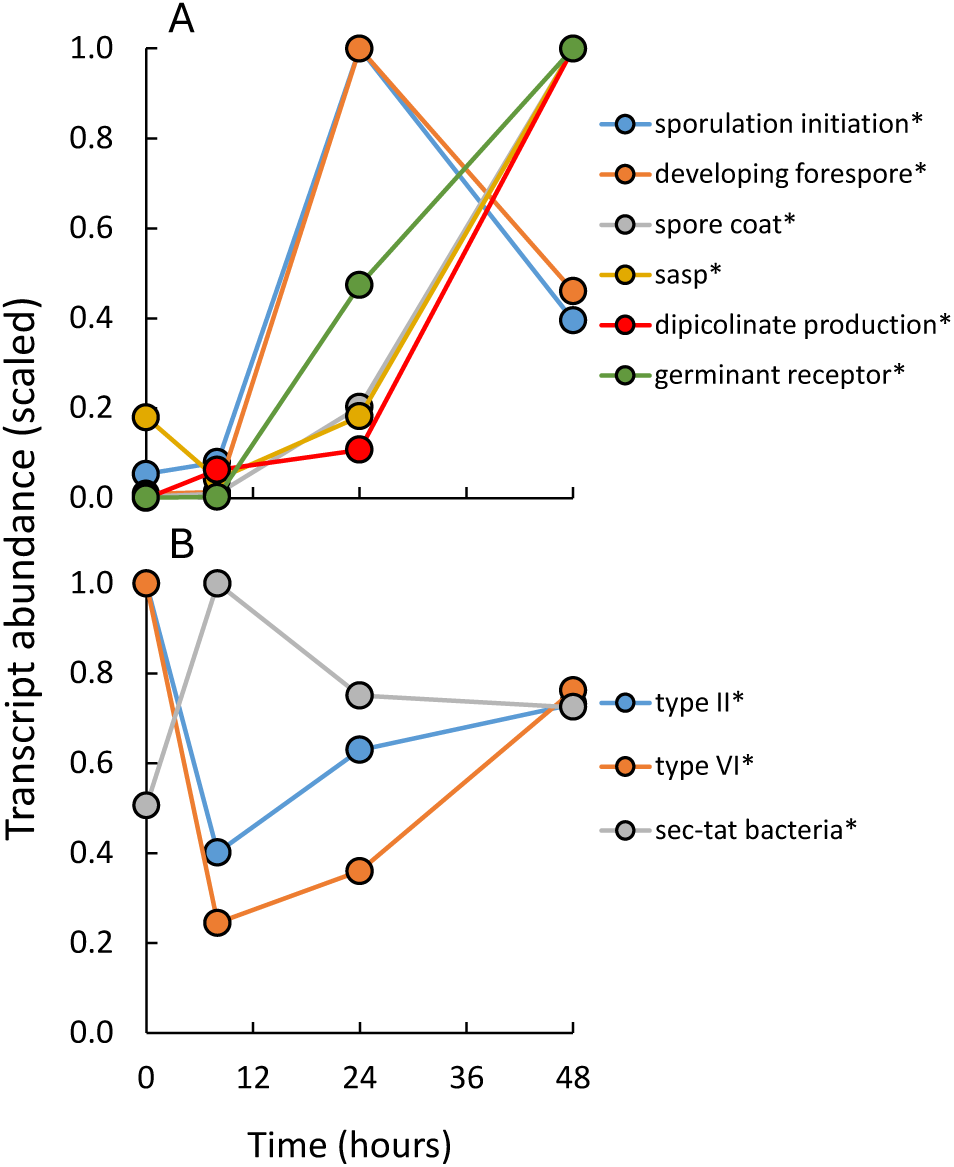
Effects of glucose addition on transcript abundances (scaled relative to maximum abundance) for genes involved in (A) sporulation initiation, developing forespore, spore coat formation, sasp proteins, dipicolinic acid production and transport, and germinant receptors, and (B) Type II and VI Secretion Systems and Sec-Tat. Sasp stands for small acid soluble proteins and Sec-Tat for Sec-dependent export system and twin arginine translocator protein export systems. * indicates statistically significant differences over time (n=4; results from multiple mean tests in Supplementary Table 4). Information on gene and transcript identity, abundances and additional statistical information is available in Supplementary Information.

### 3.8 Protein Secretion Systems

Although most proteins have a role in cytoplasm, about 30% of proteins need to be exported across the cytoplasmic membrane (Widdick et al., 2006). Several protein secretion systems have been identified (Green and Mecsas, 2016; Tsirigotaki et al., 2017). Most proteins with housekeeping functions are transported by Sec-Tat systems. Additional protein export systems include Type I-Type IX Secretion Systems. We hypothesized that after glucose addition, secretion systems with general function (Sec-Tat) would increase, while transcription of other secretion systems would decrease. In this experiment, most transcripts were related to Sec-Tat, Type II, or Type VI protein Secretion Systems (Supplementary Fig. 7B). Eight hours after glucose addition, transcript abundances for the Sec-Tat secretion system were significantly higher, while those for Type II and VI Secretion Systems significantly decreased (Fig. 6B, Supplementary Fig. 7B; Supplementary Table 4). At 24 and 48h, these trends were reversed. The increased abundance of the SEC-TAT transport system was likely associated with rapid biosynthesis, cell enlargement and growth, while that of Type VI Secretion System may be associated with bacterial predation.

## 4. Discussion

### 4.1 Feast

In response to a C addition, microbes become “activated” (Blagodatskaya and Kuzyakov, 2013; Loeppmann et al., 2020). Activation of the microbial community after C addition exhibits a general pattern of lag-phase followed by (exponential) growth. Lag phase in pure culture is defined as a period of increased metabolic activity before the first cell division following an upshift in substrate availability (Bertrand, 2019). Changes in activity in response to increased substrate availability are rapid, requiring only seconds for growing cells (Taymaz-Nikerel et al., 2011; Taymaz-Nikerel et al., 2013) or minutes to hours for dormant spores (Sinai et al., 2015; Bobek et al., 2017; Riley et al., 2020; Zhou et al., 2022). Biosynthesis of nucleotides, amino acids, lipids, and transcription and translation are all upregulated early in the lag-phase for germinating spores (Keijser et al., 2007; Sinai et al., 2015; Swarge et al., 2020) and for stationary-phase *Salmonella* cells on fresh medium (Rolfe et al., 2012). Stressful conditions cause an extended lag-phase (Hamill et al., 2020).

The length of the lag-phase in soil ecosystems is most often quantified using respiration measurements (Anderson and Domsch, 1985b; Blagodatskaya et al., 2007; Loeppmann et al., 2020) and can be anywhere from 4-18h (Blagodatskaya et al., 2007; Reischke et al., 2014; Reischke et al., 2015; Loeppmann et al., 2020). In the first studies of lag-phase in soil ecosystems, microbial growth was assumed to be absent (Anderson and Domsch, 1985b). However, in more recent investigations, microbial population growth was observed even during the lag phase (Anderson and Martens, 2013; Reischke et al., 2014; Reischke et al., 2015; Nicola and Bååth, 2019). Replication of microbes is also demonstrated in unamended bulk soil by labeling DNA with ^18^O-labeled water (Schwartz et al., 2007, Hungate et al., 2015, Metze et al 2023; Purcell et al., 2023). Comparing lag-phase in pure culture with that in soil ecosystems should be done with caution. In our experiment, we did not observe a lag-phase (Chuckran et al., 2021; Supplementay Fig. 1).

In pure culture experiments, increased substrate availability enhanced transcription and translation of genes associated with energy production and biosynthesis of all cellular compounds (e.g., Chang et al., 2002; Koburger et al., 2005; Voigt et al., 2007; Lemuth et al., 2008; Jozefczuk et al., 2010; Yamamoto et al., 2014; Irving et al., 2021). We found similar changes in this soil community: higher transcript abundances for RNA polymerase and ribosomal proteins, biosynthesis of amino acids, lipids, purine nucleotides (but not pyrimidine), cell wall synthesis, and energy production 8h after glucose addition. However, transcript abundances for degradative processes decreased temporarily (Figs. 2). Additionally, we found increased transcript abundance for regulators of the exponential phase (Fig. 5A), ribosome assembly (Fig. 5B), and Sec-Tat protein export systems (Fig. 6B) 8h after glucose addition. These transcriptional responses suggest a general upregulation of metabolism, likely resulting in enhanced replication and population expansion. This general activation does not resemble a community response where glucose is simply taken up and stored as reserve material or wasted through excess respiration (Sinsabaugh et al., 2013; Dijkstra et al., 2015; Manzoni et al 2021; Mason-Jones et al., 2021).

### 4.2 Famine

We observed the first signs of starvation 24h after glucose addition. Peak respiration occurred around 18h after glucose addition (Supplementary Fig. 1; Chuckran et al., 2021), suggesting that the transition to starvation may have occurred a few hours earlier. Evidence for starvation included (1) a reduced abundance of transcripts for regulatory genes associated with the (exponential) growth phase and increased abundance of transcripts of regulatory genes associated with stationary phase (Fig. 5A); (2) increased transcripts for early and late sporulation genes (Fig. 6A); (3) a reduction of transcripts for ribosomal proteins (Fig. 2A), assembly, and maturation (Fig. 5B) and increased abundance of transcripts associated with ribosome inactivation and hibernation (Figs. 5C); (4) an upregulation of genes associated with the glyoxylate cycle (Fig. 1A), often observed at the start of the stationary phase in pure culture (Voigt et al., 2007; Handtke et al., 2018); (5) a rebound of transcripts for degradative processes, such as nucleotide and lipid degradation (Fig. 2B); and (6) increased transcripts associated with gluconeogenesis (Fig. 2C). Cells under starvation undergo large changes in transcription in preparation for long-term survival and dormancy (Lennon and Jones, 2011; Hoehler and Jorgensen, 2013; Rittershaus et al., 2013; Wang et al., 2014; Joergensen and Wichern, 2018). These changes cause strong reductions of substrate consumption and energy demand (Pirt, 1965; Price and Sowers, 2004; Hagerty et al., 2018; Dijkstra et al., 2022), and may include (endo, exo) spore formation, cannibalism, altruistic cell death, and induction of competence (Arnaouteli et al., 2021; Qin et al., 2022). The increased ribosome hibernation, presence of stationary phase regulators, and sporulation are unlikely to be caused by the N limitation, as N-limitation was already apparent 8h after glucose addition, when transcripts for the exponential growth phase were still abundant.

Many of the regulators of exponential and stationary phase are highly species specific and therefore reflect the physiological changes for a segment of the soil community. For example, the exponential growth regulator *fis* is mostly restricted to Gammaproteobacteria, while genes for endospore formation are representative for Bacillota. Therefore, it remains unclear how much of the community transitioned to the stationary phase. Alternative responses to starvation, including utilization of alternative substrates such as acetate or prolonged zero-growth metabolism with potentially high maintenance energy demand should be further investigated.

At the start of the stationary phase (around 24h), microbial activity was still high (peak respiration 18-20h; Chuckran et al., 2021, Supplementary Fig. 1). This suggests that much of the glucose-induced respiration took place after start of the stationary phase and depletion of most of the glucose substrate (as evidenced by the low DOC content at 24h; Supplementary Fig. 1; Chuckran et al 2021). Assuming that microbial biomass was produced from the glucose substrate at relatively high efficiency (Dijkstra et al., 2011; Frey et al., 2013; Hagerty et al., 2014; Dijkstra et al., 2015, Dijkstra et al., 2022), it subsequently became susceptible to degradation via the soil food web (secondary consumption; Hagerty et al., 2014; Liang et al., 2020), explaining the large microbial activity after 24h.

### 4.3 Nutrient Limitation

Resource limitations in general, and N limitation specifically, play important roles in our understanding of C and nutrient cycles in soil ecology and ecosystem science. The soil community showed signs of N limitation 8h and 24h after glucose addition as evidenced by a rise in transcript levels for genes associated with N uptake, assimilation, and N-stress regulation (Fig. 3; Chuckran et al., 2021). Inorganic P-transporters likewise increased at 8h, albeit less than was observed for N transporters, while transcripts for phosphatases (acid, alkaline and organic-P) decreased. N limitation during rapid biosynthesis and growth makes primary consumers dependent on N from existing soil organic matter (priming; Zhang et al., 2024) and release of nutrients by the soil food web (Clarholm, 1985), likely resulting in a strong top-down trophic control over microbial productivity. The ability to identify nutrient limitations (and other stress responses) in soil communities by increased transcript abundances for functional and regulatory genes has the potential to rapidly increase our understanding of resource limitations of the soil C and nutrient cycles in managed and unmanaged ecosystems.

### 4.4 Substrate Utilization, Anabolism, Catabolism, and Biochemical Efficiency

Shortly after addition, glucose influenced the direction of C flow through the central C metabolic network, as evidenced by increased transcripts for phosphoenolpyruvate and pyruvate carboxylate 8h after glucose addition (Fig. 2C). Transcription of these enzymes is activated by glucose and suppressed in its absence (Sauer and Eikmanns, 2005; Koendjbiharie et al., 2020). However, even at 8h, gluconeogenic enzymes were present. This may be interpreted as a continued activity of gluconeogenesis and utilization of small organic C compounds or it may indicate glucose consumption using reversible gluconeogenic reactions made possible by enzymes such as Pck, Oad and MaeA (Fig. 2C). These results suggest we may be able to identify where and when microbes have access to carbohydrates in undisturbed soils. Similarly, rapid induction of transcripts associated with the glyoxylate bypass suggests that acetate was used as a substrate after glucose started to deplete (24-48h).

Before glucose was added, after a two-week preincubation, transcripts for energy production and amino acid, lipid, nucleotide, and cell wall biosynthesis were all present (Figs. 1, 2). After glucose addition, transcript abundances for these processes increased, but about in the same proportion. The transcript abundance for biosynthesis, as a proportion of the sum of transcripts for biosynthesis plus energy production (electron transport chain and central C metabolism), was 0.48 before glucose addition (Supplementary Table 4), when glucose was not a major substrate as evidenced by 1) high abundance of transcripts for genes associated with degradation suggesting substrates other than carbohydrates were being used for energy and biosynthesis (Fig. 2B); and 2) low abundance of transcripts of phosphoenolpyruvate carboxylase and pyruvate carboxylase (Fig. 2C; Sauer and Eikmanns, 2005; Koendjbiharie et al., 2020) suggesting gluconeogenesis. However, this balance between transcripts for biosynthesis and energy production showed only small though significant changes over time (proportion transcripts for biosynthesis 0.48, 0.52, 0.53 and 0.50 for t=0, 8, 24, and 48h respectively – Supplementary Table 4). These observations suggest that active biosynthesis was taking place before *and* after glucose addition, while maintaining a strong coupling between anabolic and catabolic processes. It may also suggest that proposed importance of storage reserves formation shortly after a glucose addition (Mason-Jones et al., 2019; Manzoni et al., 2021; Mason-Jones et al., 2021), potentially leading to an artificially high CUE (Hill et al., 2008; Sinsabaugh et al., 2013), may be overestimated. The latter was also suggested by a metabolic flux analysis using position-specifically labeled glucose isotopomers (Dijkstra et al., 2015). Similarly, at least at the level of transcription, there is no evidence in the transcriptome for an enhanced ‘wasteful’ respiration after glucose addition as suggested by various reviews on C use efficiency (Sinsabaugh et al., 2013; Manzoni et al., 2018; Geyer et al., 2016).

### 4.5 Biosynthesis, DNA Replication, and Cell Division

Population growth requires biosynthesis, energy production, cell enlargement, DNA replication (Morcinek-Orłowska et al., 2019), chromosome separation (Wang and Rudner, 2014), and cell division (Egan and Vollmer, 2013). Although these processes are separated in time in the cell cycle, we expect that, for an unamended soil, all processes occur simultaneously, albeit in different cells. In contrast, a pulse of exudates or added litter may temporarily ‘synchronize’ these processes.

Although we observed increases in transcripts for biosynthesis 8h after glucose addition, there were some exceptions. Biosynthesis of main lipid intermediates and products peaked 24h after glucose addition (Fig. 1, 2). This late response was also observed for transcripts of triacylglycerol and glycolipid biosynthesis genes (Supplementary Table 4). This suggests that these important energy storage and membrane compounds continued to be produced during the early stationary phase (Alvarez and Steinbuchel, 2002; Sohlenkamp and Geiger, 2015). Similarly, amino sugar and EPS production showed late peaks (24 h) in transcription (Fig. 2), suggesting a slower response of cell wall biosynthesis relative to, for example, amino acid biosynthesis. Transcript abundances for purine biosynthesis increased more than for pyrimidine synthesis, likely related to the larger role of adenine and guanine in cell metabolism (ATP, GTP, FAD, Coenzyme A, NAD(P)H in addition to RNA and DNA) compared to pyrimidine nucleosides. Similar to pyrimidines, DNA polymerase did not change significantly over time (Fig. 4A), while transcription of genes associated with deoxynucleotide production (ribonucleotide reduction; Fig. 1E) and cell division (Fig. 4B, C) was temporarily decreased 8h after glucose addition.

These results suggest that, in response to glucose addition, transcription, translation, biosynthesis and energy production increased more than cell wall production, DNA replication, and especially cell division (Table 1). The temporal disconnect between biosynthesis, DNA replication, and cell division is expected to result in increased cell size and higher DNA content per cell, possibly associated with multifork DNA replication (Trojanowski et al., 2017; Morcinek-Orłowska et al., 2019) or polyploidy (as found in aerial mycelium of Actinomycetota before spore formation; Grantcharova et al., 2005). Cell division seems to continue into the stationary phase, possibly associated with reductive cell division and spore formation (Grantcharova et al., 2005). Knowledge of the timing and sequence of these events may aid in interpreting dynamics of isotope incorporation into microbial biomass, fatty acid composition, and DNA (Gunina et al., 2014; Hungate et al., 2015; Malik et al., 2015; Müller et al 2017; Čapek et al., 2023), especially after synchronizing disturbances.

**Table 1.** Overview of changes in transcriptional abundances compared to 0h (before glucose addition).

| Process | 8h | 24h | 48h |
| --- | --- | --- | --- |
| Gluconeogenesis | ↓ | ↑ | ↑ |
| Energy | ↑ | ↓ | ↓ |
| Biosynthesis | ↑ | ↓ | ↓ |
| Cell wall production | ↑ | ↓ | ↓ |
| Ribosomes | ↑ | ↓ | ↓ |
| Degradation | ↓ | ↑ | ↑ |
| DNA polymerase | - | - | - |
| Cell division | ↓ | ↑ | ↑ |
| Stat. phase and sporulation | ↓ | ↑ | ↑ |
| N limitation | ↑ | ↑ | ↓ |

### 4.6 Secondary consumers

Secondary consumers, including predators, viruses, and scavengers, consume living and dead biomass and biofilm deposits. Also included in this category are cannibalistic bacteria, a common phenomenon during stationary phase (González-Pastor, 2011; Gophna, 2022), and utilization of internal reserves (Manzoni et al., 2021; Mason-Jones et al., 2022). The dynamics of secondary consumption in relation to primary consumption requires more study (Geisen et al., 2018; Trubl et al., 2018; Hungate et al., 2021; Petters et al., 2021; Bethany et al., 2022; Sokol et al., 2022; Dahl et al., 2023; Nicolas et al., 2023). The short burst of biomass production likely represented many new opportunities for secondary consumers, thereby explaining the continued CO_2_ production during the stationary phase (Supplementary Fig. 1).

Some progress has been made with culture studies of predator-prey interactions (Karunker et al., 2013; Laloux, 2020; Shen et al., 2021; Pérez et al., 2022; Thiery et al., 2022; Soto et al., 2023) and identification of genes associated with bacterial predators (Pasternak et al., 2013; Russell et al., 2014; Sgro et al., 2019; Hernandez et al., 2020; Gallegos-Monterrosa and Coulthurst, 2021; Thiery et al., 2022). In this study, we hypothesized that the expression of genes of Type VI Secretion System was associated with bacterial predators (Russell et al., 2014; Hernandez et al., 2020; Gallegos-Monterrosa and Coulthurst, 2021). The transcript abundance for the Type VI Secretion System genes was temporarily reduced after glucose addition (8h; Fig. 6B, Supplementary Fig. 7B), before significantly increasing at 24 and 48h, suggesting that the predator’s growth lagged the prey response by hours.

### 4.7 If Microbes are Fast, is Soil Organic Matter Formation Fast?

One main finding of this study was the rapid progression of the soil microbiome through growth and starvation following the addition of labile carbon. The initial upregulation of transcription of biosynthesis and energy production was quickly followed by transcriptional patterns typical of the stationary phase and the onset of secondary substrate consumption. This observation underscores the importance of recent findings that mineral associated organic matter is highly dynamic and contains recent C inputs (Neurath et al., 2021; Jilling et al., 2025; Sokol et al., 2025). Our study suggests that the time lag between primary and secondary consumption in response to a labile C addition may be short but could offer a window in which to stimulate soil organic matter formation processes. Defining conditions or treatments that benefit primary consumers, while reducing activity of secondary consumers, may provide a new approach for managing soil organic matter formation.

### 4.8 The Case for Metatranscriptomes

Advances in the field of metatranscriptomics have greatly increased the amount of information on soil microbial metabolism (Bei et al., 2019; Johnston et al., 2019; Albright et al., 2020; Chuckran et al., 2020; Malik et al., 2020; Nuccio et al., 2020; Chuckran et al., 2021; Kroeger et al., 2021; D’Alò et al., 2022; Söllinger et al., 2022; Guerreiro et al., 2023; Tartaglia et al., 2023), allowing us to explore new angles on general themes in soil ecology such as resource utilization and limitations, growth and efficiency, stress responses and regulation, decomposition of recalcitrant compounds, and the ecophysiology underlying respiration dynamics. It also enables us to test generally accepted theories in soil ecology with *in situ* observations. Of necessity, this will be a drawn-out process, requiring extensive knowledge of microbial metabolism at the cellular, biochemical, and molecular level. Metatranscriptome analysis also brings its own challenges: a high abundance of transcripts does not by definition mean high protein production or microbial activity (but see Balakrishnan et al., 2022). In defense of (meta)transcriptome analysis, we posit that its value for pure culture studies has been extensively proven, and thus that it, hopefully, will be equally useful in improving our understanding of microbial ecophysiology in complex soil communities.

This study summarized community-level transcript abundances per pathway and function across a large range of processes. This does not mean that each gene in a pathway or function necessarily responds in a similar way. Some genes may be associated with portions of the community that have a different response to glucose. Moreover, flux control over pathway activity may be dispersed over many reactions or confined to key steps (Grüning et al., 2025). Untangling these complex patterns is left for future studies. The overall pattern across the 1938 genes we surveyed, encompassing biosynthesis, energy production, regulatory mechanisms, and behavioral responses such as sporulation and ribosome (in)activation, together tells a story of a rapid microbiome response to a resource pulse, with microbes first entering an active phase, but quickly initiating the stationary phase when confronted with starvation (Table 1).

## Acknowledgements

This work was supported by funding from the USDA National Institute of Food and Agriculture Foundational Program (award #2017–67019-26396, #2024-67019-42340) and the U.S. Department of Energy, Office of Biological and Environmental Research, Genomic Science Program LLNL ‘Microbes Persist’ Soil Microbiome Scientific Focus Area (award #SCW1632). Work conducted by the U.S. Department of Energy Joint Genome Institute, a DOE Office of Science User Facility, was supported under contract number DE-AC02-05CH11231. Work at Lawrence Livermore National Laboratory was conducted under the auspices of the U.S. DOE under contract DE-AC52-07NA27344.

