## Supplementary Information 1 for "Rapid Changes in Transcription During a Feast-Famine Event"

**Supplemental Fig. 1**. Dynamics of (A) $CO_2$ evolution, and (B) dissolved organic C, (C) ammonium, and (D) nitrate concentrations after a glucose addition in a West Virginia agricultural soil (from Chuckran et al. 2021). $CO_2$ evolution rate was calculated as the derivative of the cumulative $CO_2$ concentrations over time using GAM fitting (R-package mgcv using REML and gratia with delta method for error propagation; Wood, 2010; Simpson, 2024). Fit was generated using k=6, select=TRUE, and gamma=1.3. The fitted line for the control treatment had an edf=0.983, a k-index = 0.48 ($p<2e^{-16}$), and an adjusted $R^2$ = 0.561, indicating no evidence of deviation from linearity. The fitted line for the plus glucose treatment had an edf = 4.66, k-index = 0.48 ($p<2e^{-16}$), and an adjusted $R^2$ = 0.993, indicating significant non-linearity. R-code for line fitting, see below.

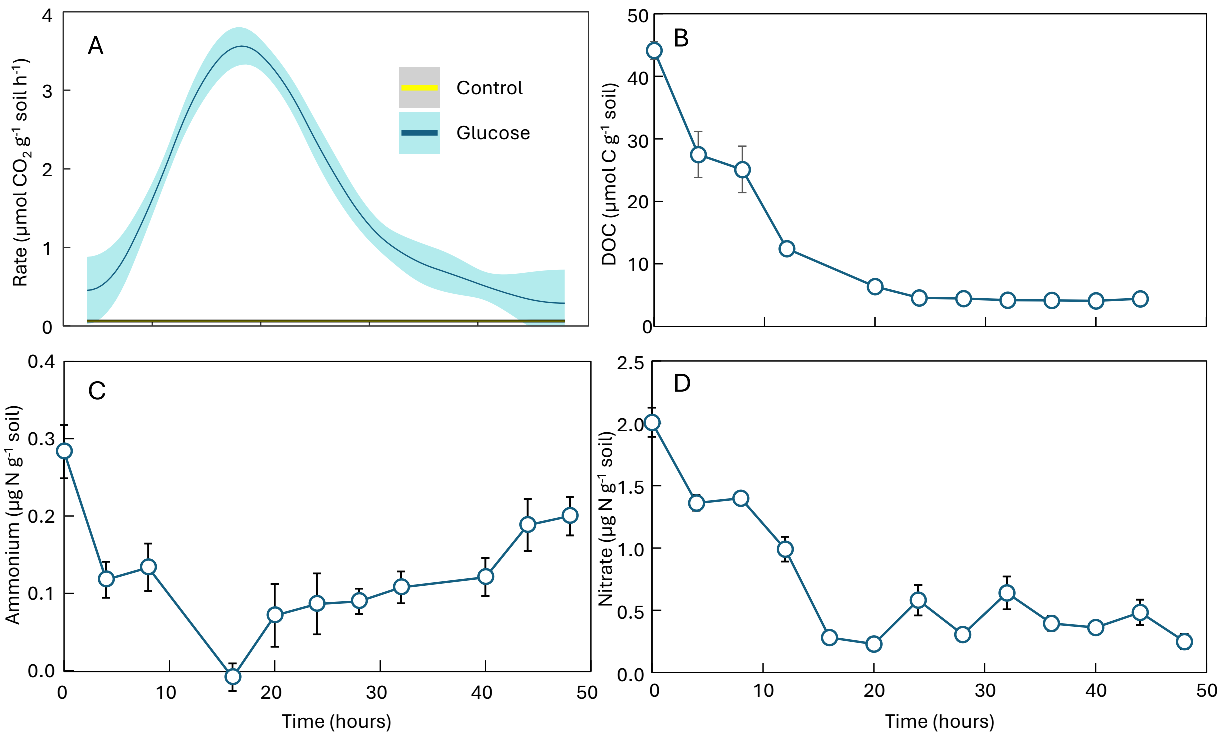


*References*


Chuckran, P.F., Fofanov, V., Hungate, B.A., Morrissey, E.M., Schwartz, E., Walkup, J., Dijkstra, P., 2021. Rapid response of nitrogen cycling gene transcription to labile carbon amendments in a soil microbial community. *mSystems* 6, e00161-00121. DOI: 10.1128/mSystems.00161-21

Simpson, G.L., 2024. gratia: An R package for exploring generalized additive models. *The Journal of Open Source Software* 9, 6962. DOI: 10.21105/joss.06962

Wood, S.N., 2010. Fast stable restricted maximum likelihood and marginal likelihood estimation of semiparametric generalized linear models. *Journal of the Royal Statistical Society Series B: Statistical Methodology* 73, 3-36. DOI: 10.1111/j.1467-9868.2010.00749.x

*R-code*

```
# Code to calculate CO₂ production rate over time from cumulative CO₂ production
library(mgcv)    # GAM fitting - general additive model
library(gratia)  # Derivatives and uncertainty using delta method
library(ggplot2)
library(dplyr)
library(readr)
# Set file location
df <- read_csv("CO2_data.csv")
# Separate treatments
Control <- df %>% filter(treatment == "C")
Glucose <- df %>% filter(treatment == "G")
# Run generalized additive model (gam)
Gam_C <- gam(CO2 ~ s(time, k = 6), data = Control, method = "REML", select = TRUE, gamma = 1.3)
Gam_G <- gam(CO2 ~ s(time, k = 6), data = Glucose, method = "REML", select = TRUE, gamma = 1.3)
summary(Gam_C)
summary(Gam_G)
# Combine results in one file for plotting
p_smooth <- bind_rows(
  fitted_values(Gam_C) %>% mutate(treatment = "Control"),
  fitted_values(Gam_G) %>% mutate(treatment = "Glucose"))
# The p_smooth file contains some counter-intuitive names. For simplicity rename
p_smooth <- p_smooth %>%
  rename(fitted = .fitted, se = .se, lower_CI = .lower_ci, upper_CI = .upper_ci)
# plotting cumulative CO2 production
ggplot(p_smooth, aes(x = time, y = fitted, color = treatment)) + geom_line(size = 1) +
  geom_ribbon(aes(ymin = lower_CI, ymax = upper_CI, fill = treatment), alpha = 0.3, color = NA) +
  labs(title = "GAM Smooths by Treatment", x = "x", y = "Fitted y") + theme_bw()
# Calculate derivatives
Deriv_C <- derivatives(Gam_C, select = "s(time)")
Deriv_G <- derivatives(Gam_G, select = "s(time)")
# Adding column with treatment names
Deriv_C$treatment <- "Control"
Deriv_G$treatment <- "Glucose"
# Combine in one data frame
Deriv_All <- bind_rows(Deriv_C, Deriv_G)
# Rename variables
Deriv_All <- Deriv_All %>%
  rename(rate = .derivative, se = .se, crit = .crit, lower_CI = .lower_ci, upper_CI = .upper_ci)
p<-ggplot(Deriv_All, aes(x = time, y = rate, color = treatment)) + geom_line(size = 1) +
  geom_ribbon(aes(ymin = lower_CI, ymax = upper_CI, fill = treatment), alpha = 0.3, color = NA) +
```

```
  geom_hline(yintercept = 0, linetype = "dashed") + labs(title = "CO2 production rates", x = "time",
    y = “rate (micromole CO2 g-1 soil h-1)”) + theme_bw()
# Save as svg plot for import in PowerPoint
ggsave("derivative_plot.svg", p)
```
