## Supplementary Information 2 for "Rapid Changes in Transcription During a Feast-Famine Event"

| Main Text | Supplementary Figures | | |
|---|---|---|---|
| Transcript abundance (most scaled) | Transcript abundance (most per million) | Gene abundance (scaled) | Gene abundance (per million) |
| Fig. 1 | Fig. 2 | Fig. 8 | Fig. 14 |
| Fig. 2 | Fig. 3 | Fig. 9 | Fig. 15 |
| Fig. 3 | Fig. 4 | Fig. 10 | Fig. 16 |
| Fig. 4 | Fig. 5 | Fig. 11 | Fig. 17 |
| Fig. 5 | Fig. 6 | Fig. 12 | Fig. 18 |
| Fig. 6 | Fig. 7 | Fig. 13 | Fig. 19 |

**Supplementary Fig. 2**. Transcript abundances (per million transcripts) for genes associated with (A) central C metabolic network, (B) electron transport chain, biosynthesis of (C) amino acids, (D) lipids, (E) nucleotides, and (F) cell wall compounds and exopolysaccharides. * indicates statistically significant differences across time. Glycol., PYR ox. stands for glycolysis and pyruvate oxidation, pw. for pathway, and eps for exopolysaccharides. Amino acid biosynthesis pathways are represented by their one-letter amino acid code. Additional information on gene and transcript identity, abundances and additional statistical information is available in Supplementary Tables.

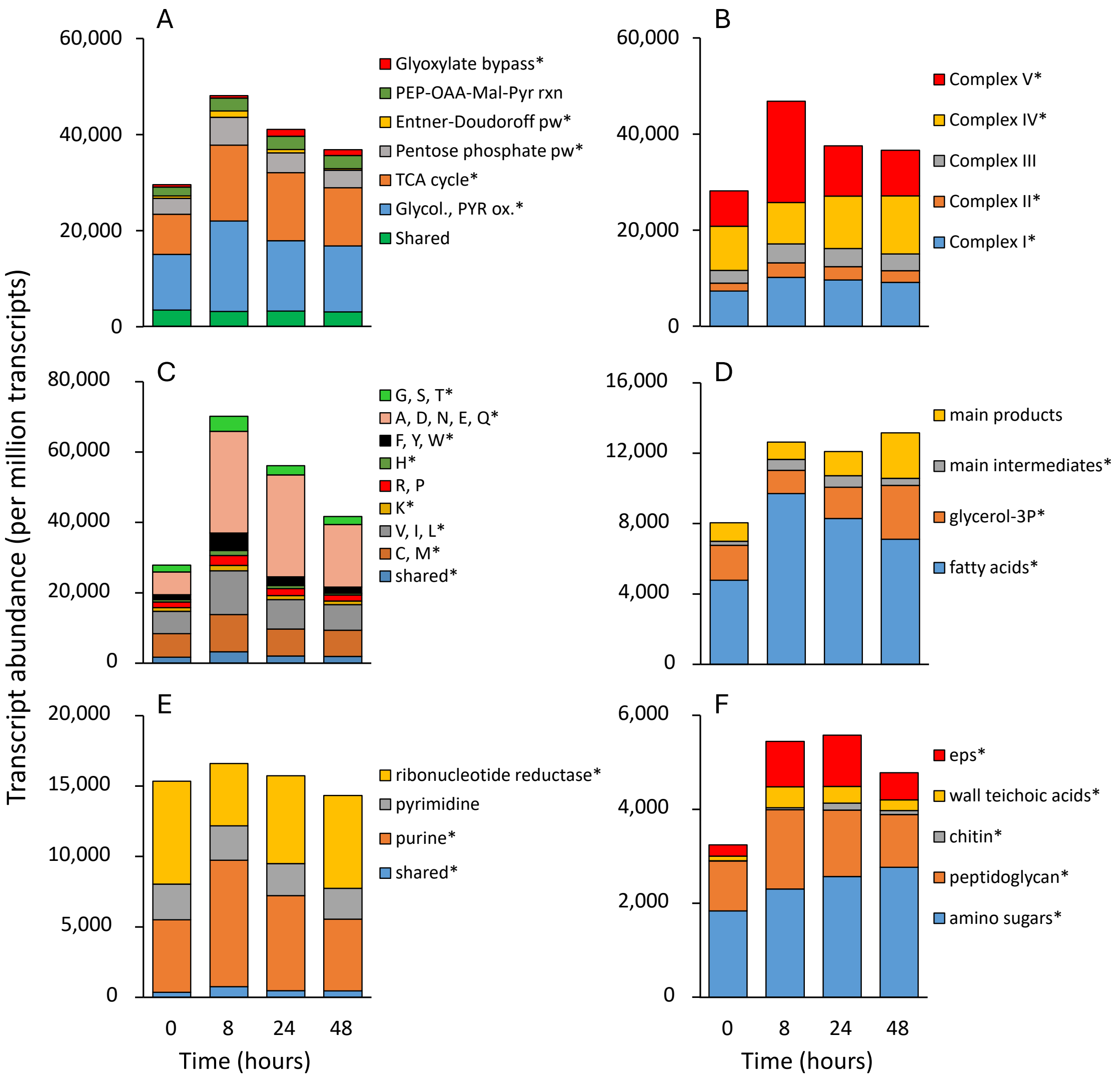

**Supplementary Fig. 3**. Transcript abundances (per million transcripts) for genes associated with (A) biosynthesis of amino acids, lipids, nucleotides, cell walls and exopolysaccharides, (B) degradation of amino acids, lipids, nucleotides, and (hetero)cyclic hydrocarbons, and (C) genes for glycolytic (pyruvate carboxylase - *pyc*, phosphoenolpyruvate carboxylase - *ppc*) and gluconeogenic (oxaloacetate decarboxylase - *oad*, (oxaloacetate-decarboxylating) malate dehydrogenase - *maeA*, and phosphoenolpyruvate carboxykinase - *pck, pepck*) reactions between PEP, pyruvate and oxaloacetate and malate. * indicates statistically significant differences across time. Additional information on gene and transcript identity, abundances and additional statistical information is available in Supplementary Tables.

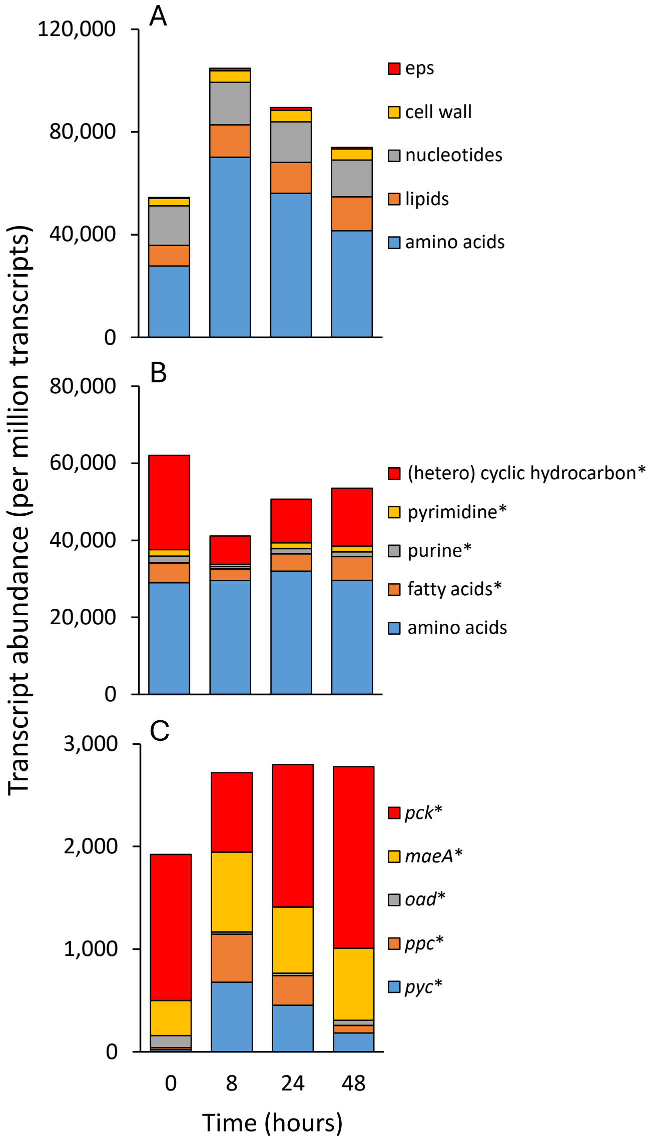

**Supplementary Fig. 4**. Transcript abundance (scaled) for (A) regulator genes associated with C (*crp*), P (*phoB-phoR*), and N stress (*ntrC-ntrB*), (B) P-transporters and phosphatases, and (C) ammonium (*amt*) and nitrate transporters, GS-GOGAT (*glnA*, *glt*) and regulatory genes (*glnK*, *glnD*). * indicates statistically significant differences across time (Table S4). Information on gene and transcript identity, abundances and additional statistical information is available in Supplementary Tables.

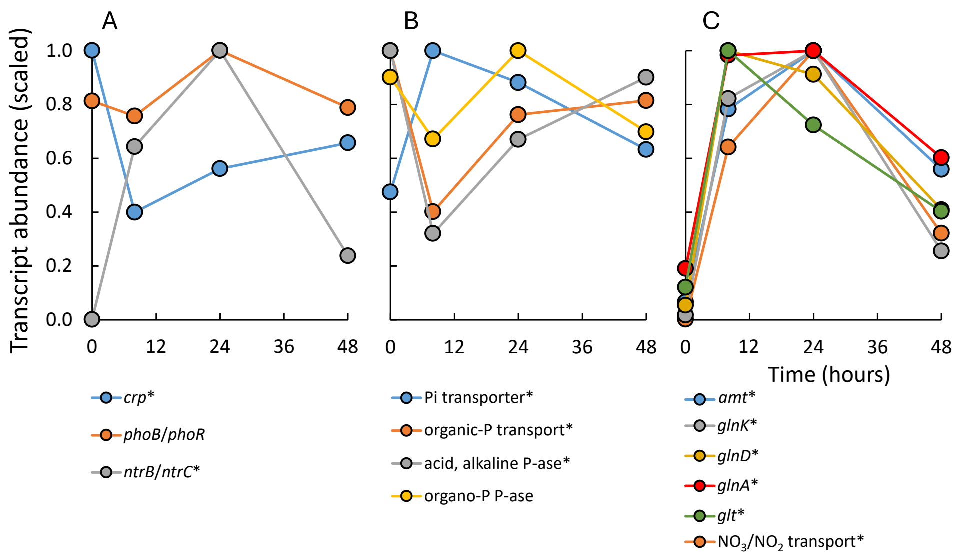

**Supplementary Fig. 5**. Transcript abundances (per million transcripts) for (A) DNA polymerase III, RNA polymerase, and ribosomal proteins, (B) genes of early stage of cell division (*ftsZ, ftsA, divIVA)* and *(*C) mid-stage of cell division (*ftsK, ftsW, ftsI, and sepF*). * indicates statistically significant differences over time. Information on gene and transcript identity, abundances and additional statistical information is available in Supplementary Tables.

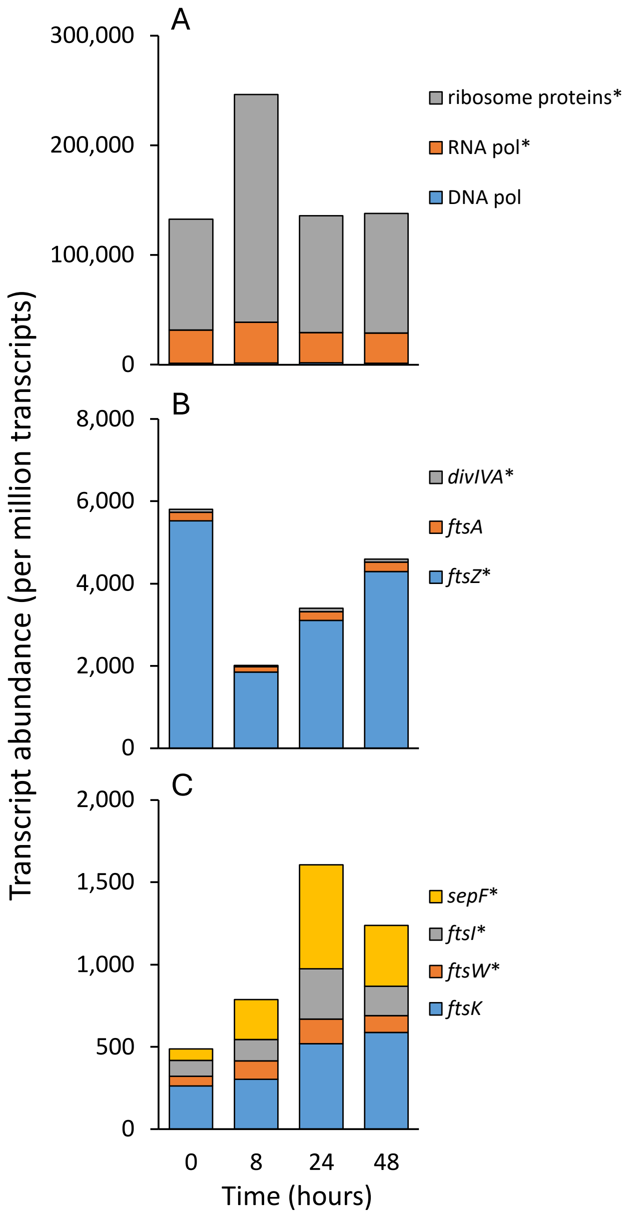

**Supplementary Fig. 6**. Transcript abundances (per million transcripts) for (A) regulators of the early exponential growth phase (*fis*), late exponential growth phase (*hns*), and early stationary phase (*rpoS*, *rsd*), (B) ribosome activation (*rbfA*, *rimP*, and *rimM*) and (C) ribosome hibernation (*rmf*, *hpf*, *ybeB*). * indicates statistically significant differences over time. Information on gene and transcript identity, abundances and additional statistical information is available in Supplementary Tables.

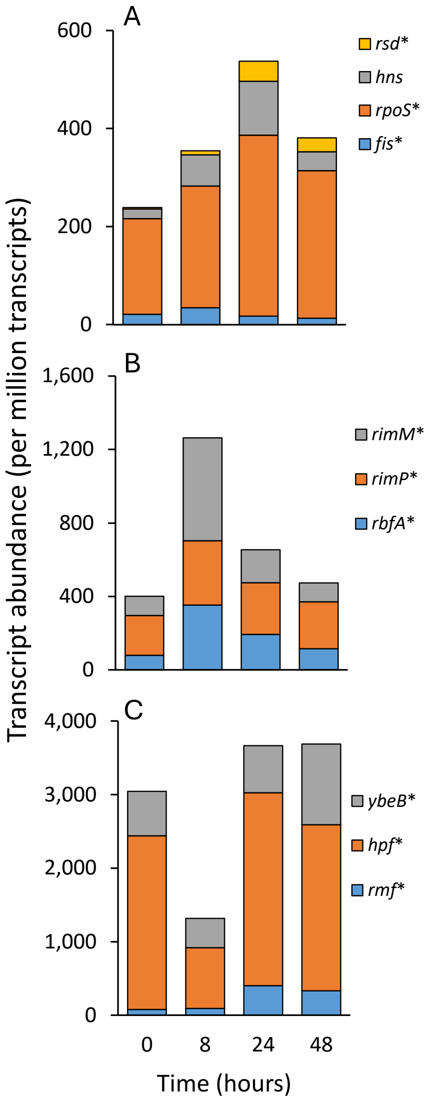

**Supplementary Fig. 7**. Transcript abundances (per million transcripts) for genes involved in (A) sporulation initiation, developing forespore, spore coat formation, sasp proteins, dipicolinic acid production and transport, and germinant receptors, and (B) Type II and VI secretion systems and Sec-Tat. Sasp stands for small acid soluble proteins and Sec-Tat for Sec-dependent export system and twin arginine translocator protein export systems. Information on gene and transcript identity, abundances and additional statistical information is available in Supplementary Tables.

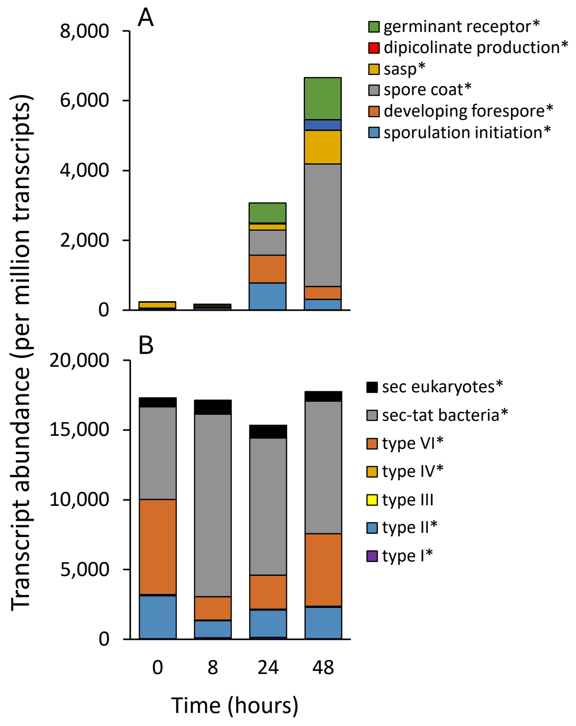

**Supplementary Fig. 8**. Gene abundances (scaled) for genes associated with (A) central C metabolic network, (B) electron transport chain, biosynthesis of (C) amino acids, (D) lipids, (E) nucleotides, and (F) cell wall compounds and exopolysaccharides. * indicates statistically significant differences across time. Glycol., PYR ox. stands for glycolysis and pyruvate oxidation, pw. for pathway, and eps for exopolysaccharides. Amino acid biosynthesis pathways are represented by their one-letter amino acid code. Additional information on gene and transcript identity, abundances and additional statistical information is available in Supplementary Tables.

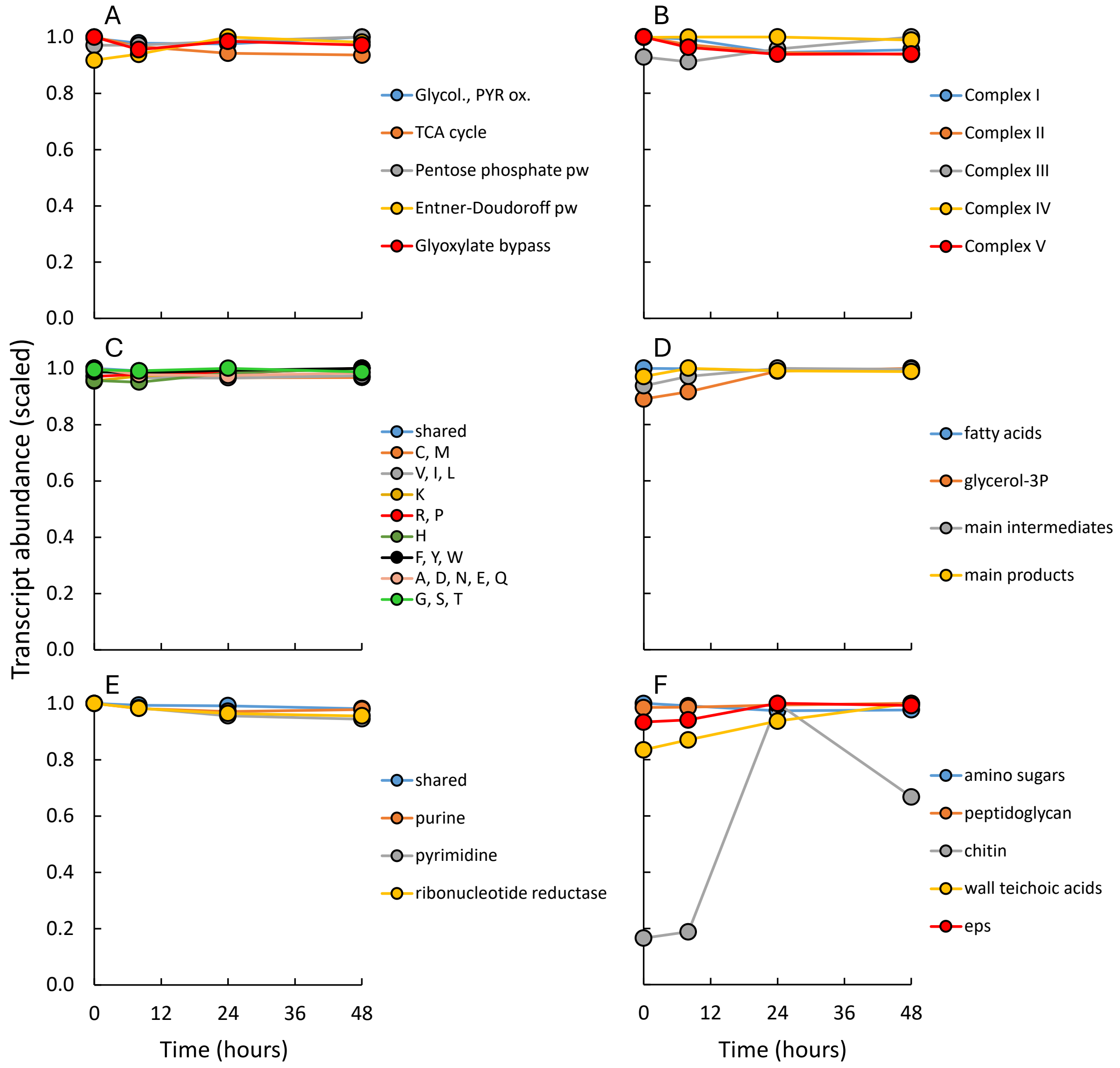

**Supplementary Fig. 9**. Gene abundances (scaled) for genes associated with (A) biosynthesis of amino acids, lipids, nucleotides, cell walls and exopolysaccharides, (B) degradation of amino acids, lipids, nucleotides, and (hetero)cyclic hydrocarbons, and (C) genes for glycolytic (pyruvate carboxylase - *pyc*, phosphoenolpyruvate carboxylase - *ppc*) and gluconeogenic (oxaloacetate decarboxylase - *oad*, (oxaloacetate-decarboxylating) malate dehydrogenase - *maeA*, and phosphoenolpyruvate carboxykinase - *pck, pepck*) reactions between PEP, pyruvate and oxaloacetate and malate. * indicates statistically significant differences across time. Additional information on gene and transcript identity, abundances and additional statistical information is available in Supplementary Tables.

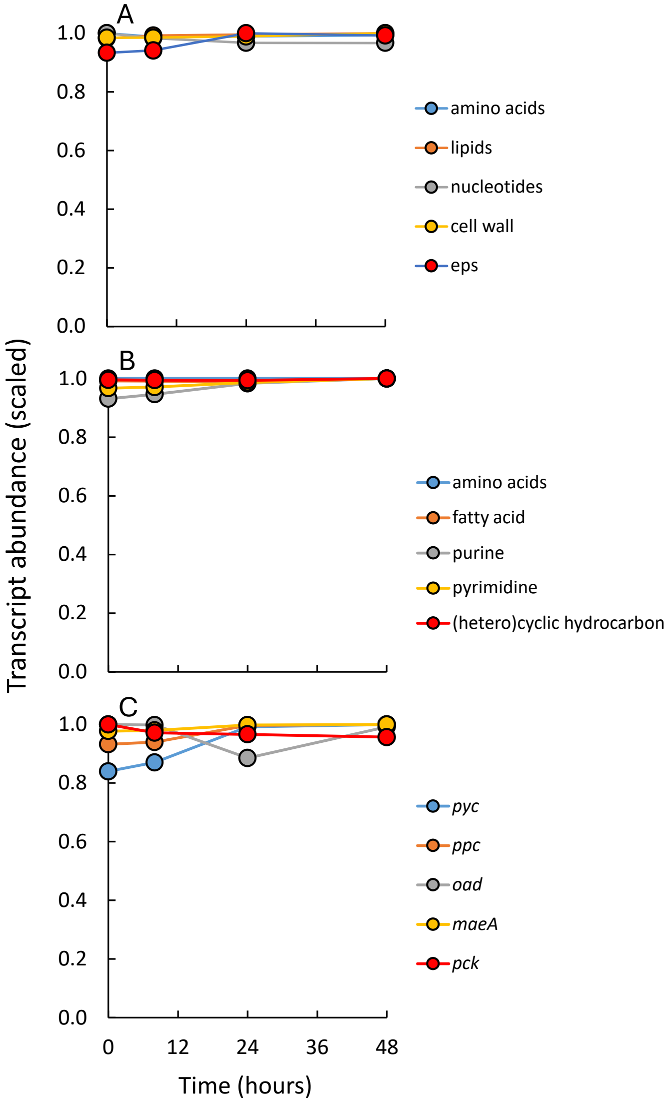

**Supplementary Fig. 10**. Gene abundance (scaled) for (A) regulator genes associated with C (*crp*), P (*phoB-phoR*), and N stress (*ntrC-ntrB*), (B) P-transporters and phosphatases, and (C) ammonium (*amt*) and nitrate transporters, GS-GOGAT (*glnA*, *glt*) and regulatory genes (*glnK*, *glnD*). * indicates statistically significant differences across time (Table S4). Information on gene and transcript identity, abundances and additional statistical information is available in Supplementary Tables.

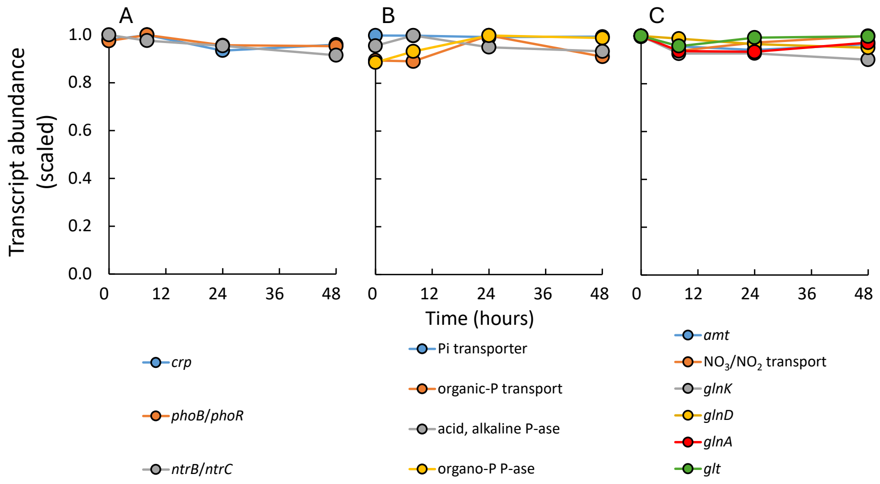

**Supplementary Fig. 11**. Gene abundances (scaled) for (A) DNA polymerase III, RNA polymerase, and ribosomal proteins, (B) genes of early stage of cell division (*ftsZ, ftsA, divIVA)* and *(*C) mid-stage of cell division (*ftsK, ftsW, ftsI, and sepF*). * indicates statistically significant differences over time. Information on gene and transcript identity, abundances and additional statistical information is available in Supplementary Tables.

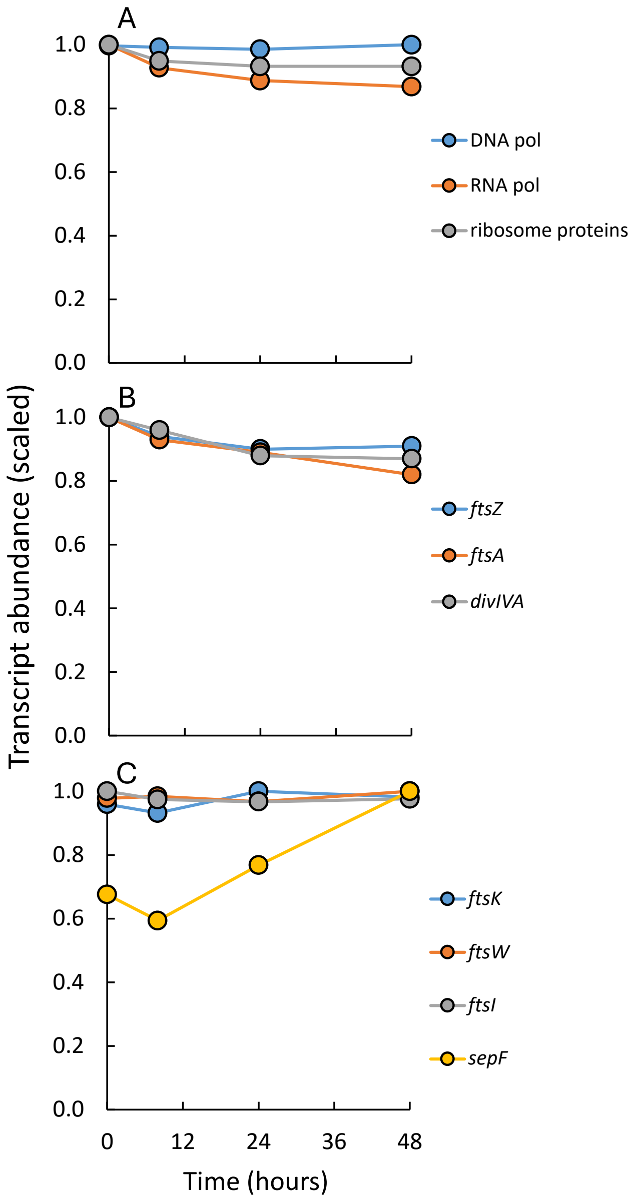

**Supplementary Fig. 12**. Gene abundances (scaled) for (A) regulators of the early exponential growth phase (*fis*), late exponential growth phase (*hns*), and early stationary phase (*rpoS*, *rsd*), (B) ribosome activation (*rbfA*, *rimP*, and *rimM*) and (C) ribosome hibernation (*rmf*, *hpf*, *ybeB*). * indicates statistically significant differences over time. Information on gene and transcript identity, abundances and additional statistical information is available in Supplementary Tables.

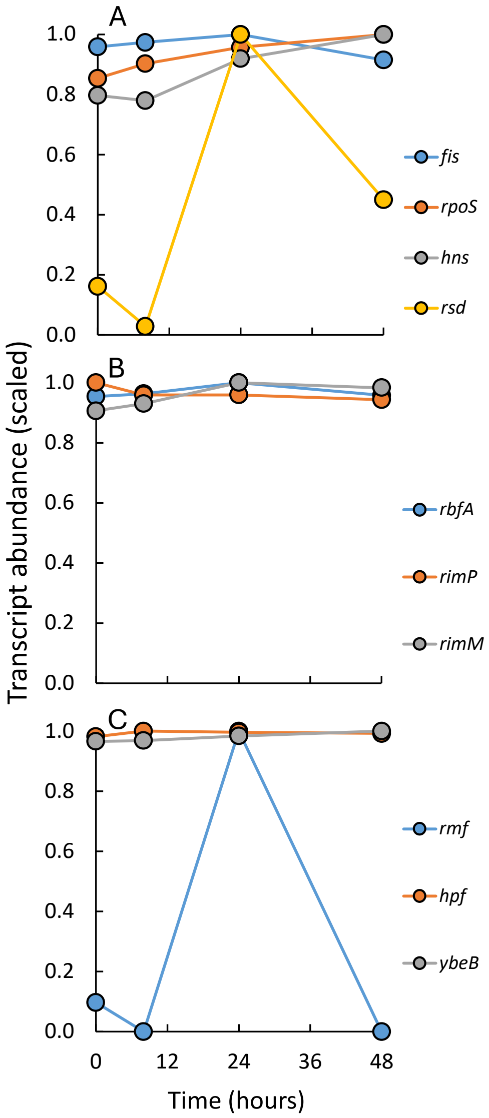

**Supplementary Fig. 13**. Gene abundances (scaled) for genes involved in (A) sporulation initiation, developing forespore, spore coat formation, sasp proteins, dipicolinic acid production and transport, and germinant receptors, and (B) Type II and VI secretion systems and Sec-Tat. Sasp stands for small acid soluble protein and Sec-Tat for Sec-dependent export system and twin arginine translocator protein export systems. Information on gene and transcript identity, abundances and additional statistical information is available in Supplementary Tables.

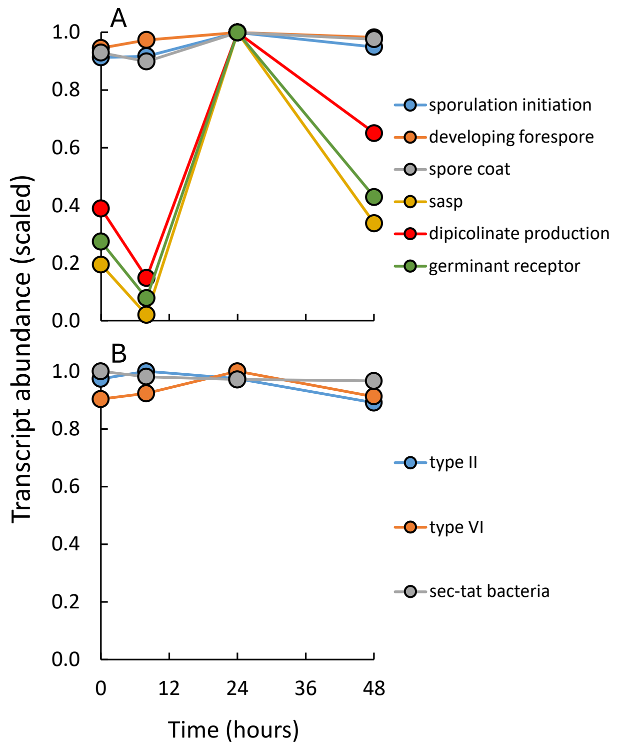

**Supplementary Fig. 14**. Gene abundances (per million genes) for genes associated with (A) central C metabolic network, (B) electron transport chain, biosynthesis of (C) amino acids, (D) lipids, (E) nucleotides, and (F) cell wall compounds and exopolysaccharides. * indicates statistically significant differences across time. Glycol., PYR ox. stands for glycolysis and pyruvate oxidation, pw. for pathway, and eps for exopolysaccharides. Amino acid biosynthesis pathways are represented by their one-letter amino acid code. Additional information on gene and transcript identity, abundances and additional statistical information is available in Supplementary Tables.

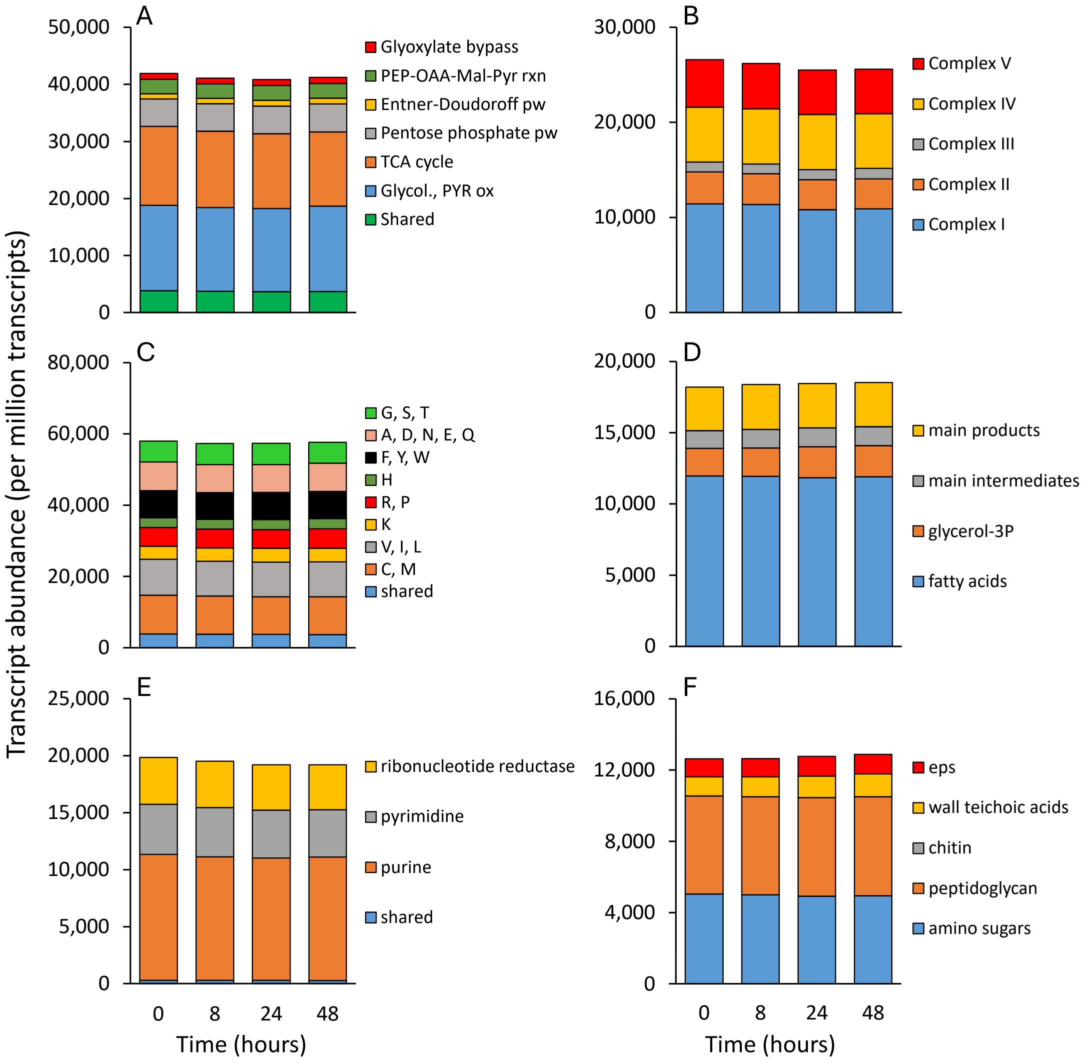

**Supplementary Fig. 15**. Gene abundances (per million genes) for genes associated with (A) biosynthesis of amino acids, lipids, nucleotides, cell walls and exopolysaccharides, (B) degradation of amino acids, lipids, nucleotides, and (hetero)cyclic hydrocarbons, and (C) genes for glycolytic (pyruvate carboxylase - *pyc*, phosphoenolpyruvate carboxylase - *ppc*) and gluconeogenic (oxaloacetate decarboxylase - *oad*, (oxaloacetate-decarboxylating) malate dehydrogenase - *maeA*, and phosphoenolpyruvate carboxykinase - *pck, pepck*) reactions between PEP, pyruvate and oxaloacetate and malate. * indicates statistically significant differences across time. Additional information on gene and transcript identity, abundances and additional statistical information is available in Supplementary Tables.

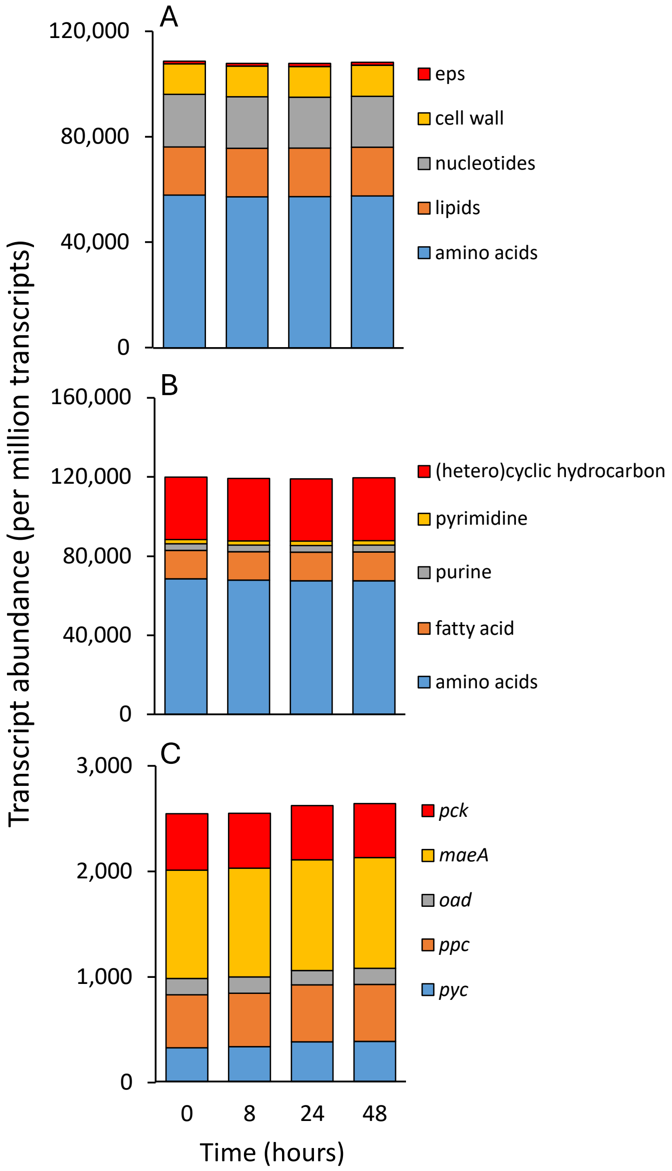

**Supplementary Fig. 16**. Gene abundance (transcripts per million genes) for (A) regulator genes associated with C (*crp*), P (*phoB-phoR*), and N stress (*ntrC-ntrB*), (B) P-transporters and phosphatases, and (C) ammonium (*amt*) and nitrate transporters, GS-GOGAT (*glnA*, *glt*) and regulatory genes (*glnK*, *glnD*). * indicates statistically significant differences across time (Table S4). Information on gene and transcript identity, abundances and additional statistical information is available in Supplementary Tables.

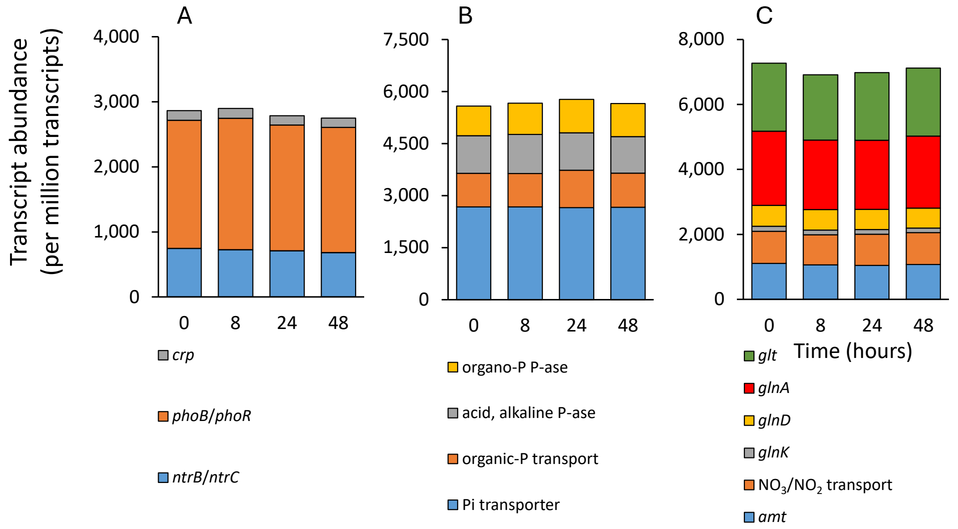

**Supplementary Fig. 17**. Gene abundances (per million genes) for (A) DNA polymerase III, RNA polymerase, and ribosomal proteins, (B) genes of early stage of cell division (*ftsZ*, *ftsA*, *divIVA*) and (C) mid-stage of cell division (*ftsK*, *ftsW*, *ftsI*, and *sepF*). * indicates statistically significant differences over time. Information on gene and transcript identity, abundances and additional statistical information is available in Supplementary Tables.

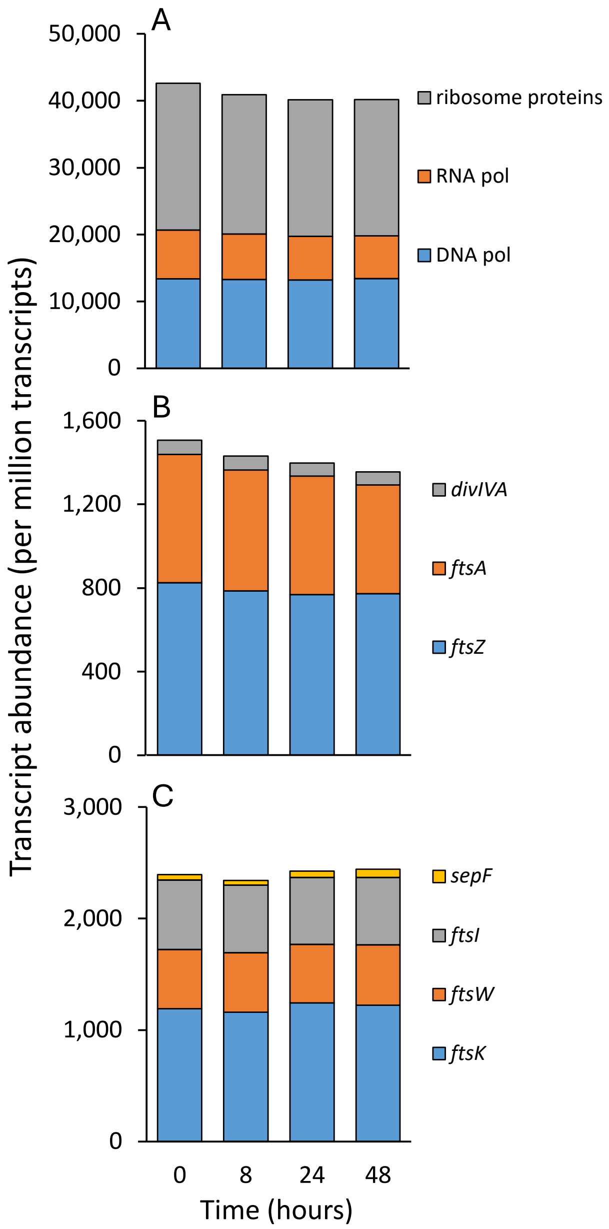

**Supplementary Fig. 18**. Gene abundances (per million genes) for (A) regulators of the early exponential growth phase (*fis*), late exponential growth phase (*hns*), and early stationary phase (*rpoS*, *rsd*), (B) ribosome activation (*rbfA*, *rimP*, and *rimM*) and (C) ribosome hibernation (*rmf*, *hpf*, *ybeB*). * indicates statistically significant differences over time. Information on gene and transcript identity, abundances and additional statistical information is available in Supplementary Tables.

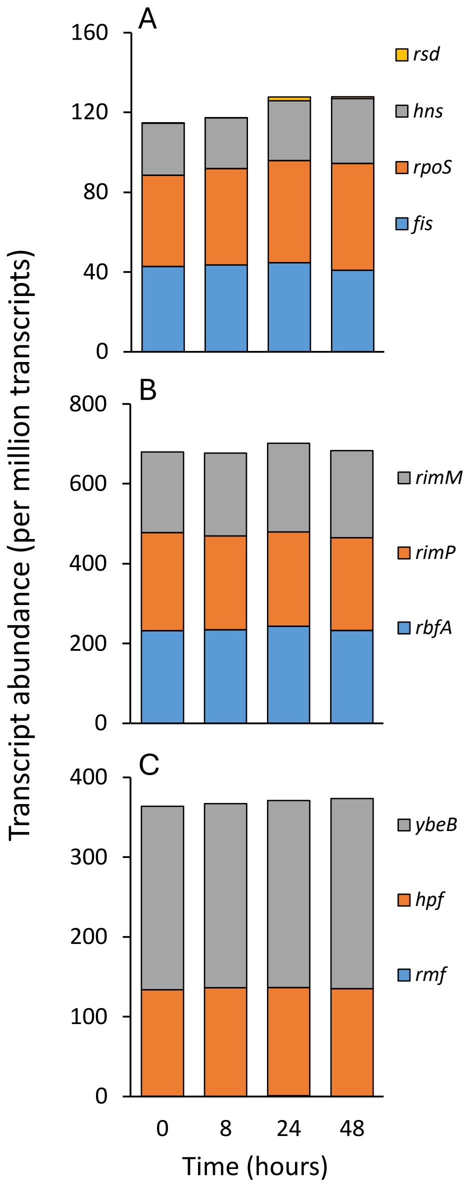

**Supplementary Fig. 19**. Gene abundances (per million genes) for genes involved in (A) sporulation initiation, developing forespore, spore coat formation, sasp proteins, dipicolinic acid production and transport, and germinant receptors, and (B) Type II and VI secretion systems and Sec-Tat. Sasp stands for small acid soluble protein and Sec-Tat for Sec-dependent export system and twin arginine translocator protein export systems. Information on gene and transcript identity, abundances and additional statistical information is available in Supplementary Tables.

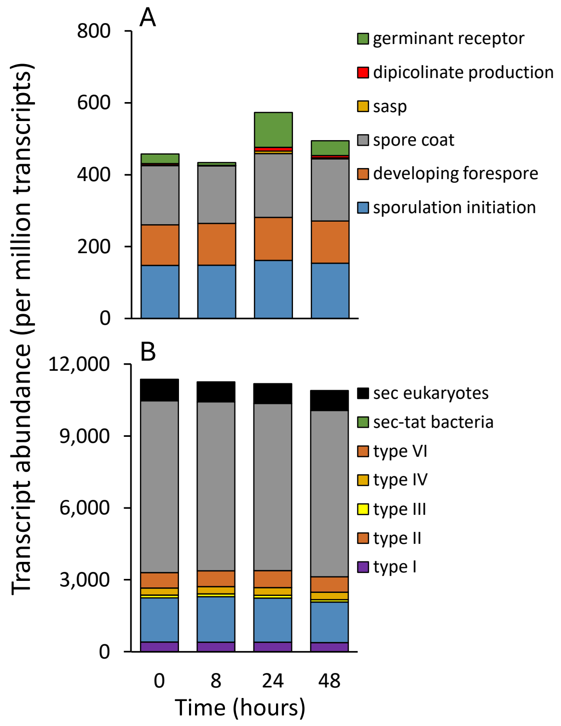
